# Altered circadian rhythm, sleep, and *rhodopsin 7*-dependent shade preference during diapause in *Drosophila melanogaster*

**DOI:** 10.1101/2024.03.09.584236

**Authors:** Geoff T. Meyerhof, Sreesankar Easwaran, Angela E. Bontempo, Craig Montell, Denise J. Montell

**Affiliations:** Department of Molecular, Cellular, and Developmental Biology and the Neuroscience Research Institute, University of California, Santa Barbara, California 93106, USA ORCID IDs: SE, 0000-0003-4676-2150, CM, 0000-0001-5637-1482, DJM, 0000-0001-8924-5925

**Keywords:** Diapause, *Drosophila*, sleep, circadian rhythm, rhodopsin

## Abstract

To survive adverse environments, many animals enter a dormant state such as hibernation, dauer, or diapause. Various *Drosophila* species undergo adult reproductive diapause in response to cool temperatures and/or short day-length. While it is known that flies are less active during diapause, an in-depth understanding of diapause effects on circadian rhythms and sleep is lacking. Here we show that, in diapause-inducing conditions, *Drosophila melanogaster* exhibit altered circadian activity profiles, including a severely reduced morning activity peak and an advanced evening activity peak. Consequently, the flies have a single activity peak at a time similar to when non-diapausing flies have a siesta. Temperatures ≤15 °C, rather than short day-length, primarily drive the behavior. At cool temperatures, flies also rapidly enter a deep sleep state that lacks the sleep cycles of flies at higher temperatures and requires particularly high levels of stimulation for arousal. Furthermore, we show that at 18–25 °C, flies prefer to siesta in the shade, a preference that is virtually eliminated at 10 °C. Resting in the shade is driven by an aversion to blue light, sensed by rhodopsin 7 (Rh7) outside of the eyes. Flies at 10 °C show neuronal markers of elevated sleep pressure, including increased expression of Bruchpilot and elevated Ca^2+^ in the R5 ellipsoid body neurons. Therefore sleep pressure might overcome blue light aversion. Thus at temperatures known to cause reproductive arrest, preserve germline stem cells, and extend lifespan, *Drosophila melanogaster* are prone to deep sleep and exhibit dramatically altered – yet rhythmic – daily activity patterns.

**Significance statement:** Climate change is impacting many animals, including insects. In diverse organisms, adverse environments trigger dormancy programs such as hibernation and diapause. Fruit flies undergo diapause to survive winter. Here we develop new methods and show that the same cool temperatures that delay fruit fly reproduction and extend lifespan, also promote deep sleep. Cool flies rapidly fall asleep and are difficult to arouse. Once awake, they immediately fall back to sleep. Whereas in warm environments, midday blue light drives flies to siesta in the shade, in cool temperatures the need to sleep overwhelms light-aversion, reducing shade preference. Animals that adjust their behavior directly to temperature, rather than day length, may be more resilient to a changing climate.

## Introduction

As the climate changes, understanding organismal responses to temperature becomes more important. Dormancy programs such as hibernation, torpor, dauer and diapause allow animals to endure adverse environments while extending lifespan and reproductive capacity (1–5). Diapause is a well-documented phenomenon in numerous insect species including *Drosophila*, as well as in animals such as killifish and the mouse, (6, 7). Diapause contributes to survival as it allows animals to halt their development and/or reproduction, and delay aging, until environmental conditions improve (8).

Various *Drosophila* species undergo adult reproductive diapause in response to cool temperatures and/or short day length (9–11). The conditions that induce diapause not only arrest growth and development, and extend lifespan, but they also alter metabolism and behavior. This is critical as survival depends on matching food intake and activity levels with energy needs. Flies in diapause reduce food intake and overall activity yet maintain high levels of circulating nutrients (12). Although initially characterized as an arrest of ovarian development at the yolk-uptake stage, diapause is now recognized to be a complex program that affects nearly every aspect of the animal’s life. However it is not known if the reduced activity in diapause represents a cool-temperature-induced immobility or an actively modified behavioral program.

Significant progress is being made to elucidate the molecular and cellular effects of diapause conditions on fly physiology, metabolism, reproduction and lifespan (12–17). For example, cool temperatures dampen activity in subsets of circadian pacemaker neurons, reducing levels of secreted neuropeptides and hormones and thereby arrest ovarian development (17, 18). So it is clear that sensing environmental conditions, primarily temperature, precedes and leads to reproductive arrest.

Many open questions remain (8). For example, an in-depth understanding of diapause effects on behavior, especially circadian rhythms and sleep characteristics, is lacking. Here we describe that at 10-15 °C, flies show a dramatically altered circadian rhythm. This effect is more dependent on temperature than photoperiod. Using a suite of behavioral assays, we show that flies maintained in cool conditions rapidly fall into deep sleep and become difficult to arouse, although their remaining activity retains rhythmicity. We show that at warm temperatures, flies prefer to siesta in the shade due to aversion of blue light, which is sensed by rhodopsin 7 (Rh7). At the same cool temperatures that induce reproductive diapause, sleep pressure overwhelms this blue light aversion, reducing the preference for sleeping in the shade. The effects of cool temperatures on circadian rhythm and sleep are independent of juvenile hormone (JH). Thus the same cool temperatures that induce reproductive arrest via decreased JH production, cause JH-independent behavioral effects.

## Results

### Diapause-inducing conditions reshape the *Drosophila melanogaster* circadian activity profile

To determine the effect of diapause-inducing conditions [10°C and 8 h L and 16 h D cycles (LD8:16)] on circadian activity, we first employed the *Drosophila* Activity Monitor (DAM) system to record the movement of flies throughout the day (19, 20). In this assay, flies are individually housed in clear glass tubes with constant access to a sucrose food source. Their activity is recorded via an infrared sensor that runs through the center of each tube. We allowed female flies to acclimatize to either 25 °C or 10 °C for ≥24 hours (h) under varying light/dark (LD) cycles, and then recorded their activities for four days. At 25 °C in LD8:16, the flies displayed prominent morning and evening activity peaks, which flanked a period of inactivity in the middle of the day known as the siesta (Fig. 1*A*, red lines) (21). In this short photoperiod condition (LD8:16), the flies’ morning activity peak preceded lights on, and their evening activity peaked at lights off. Thus, the majority of their activity occurred at night (Fig. 1*A*), consistent with a recent report (22).

**Fig. 1.**
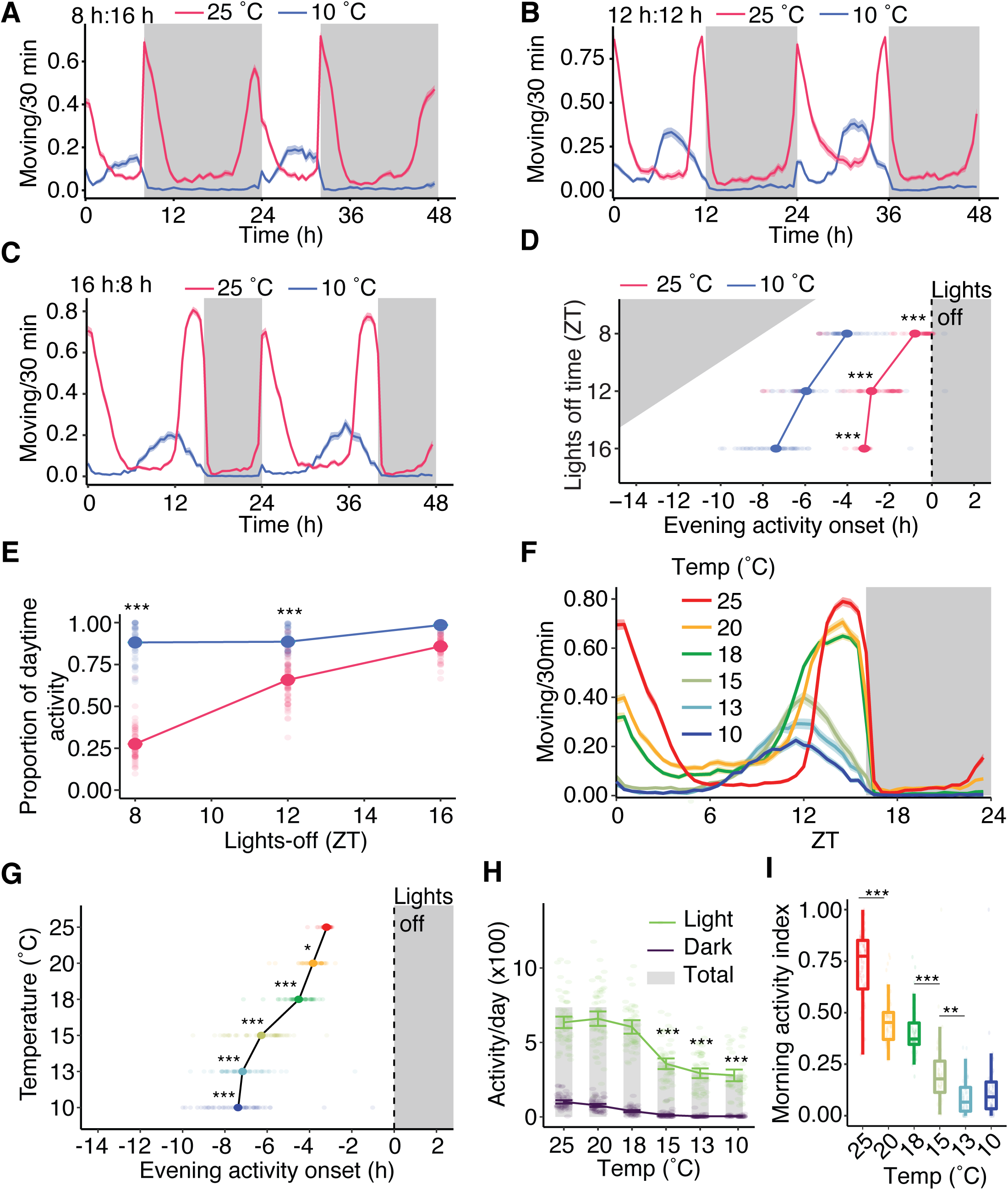
Diapause shifts activity pattern from crepuscular to peak midday activity. (*A*) 48-hour average actogram of flies during 8 hr light:16 h dark cycles housed at 25 °C (red) or 10 °C (blue). n = 63 flies at 25 °C and 46 flies at 10 °C.(*B*) 48-hour average actogram of flies exposed to 12 hr light:12 hr dark cycles at 25 °C (red) or 10 °C (blue). n = 74 flies at 25 °C and 55 flies at 10 °C. (*C*) 48-hour average actogram of flies under 16 hr light:8 hr dark cycles housed under 25 °C (red) or 10 °C (blue). n = 64 flies at 25 °C and 51 flies at 10 °C. (*D*) Time of evening activity peak onset versus lights-off time (i.e., photoperiod) from panels *A*, *B*, and *C*. Two-way ANOVA with factors of temperature, photoperiod, and temperature:photoperiod interaction. Individual group differences calculated using the Tukey HSD test. (*E*) Proportion of activity occurring during daytime from *A, B, and C* Two-way ANOVA with factors of light,temperature, and light:temperature interaction. Individual differences calculated using the Tukey HSD test. (*F*) Effect of temperature on the circadian activity profile of flies housed at 25 °, 20 °, 18 °, 15 °, 13 °, and 10 °C. n=51–64 flies/temperature. (*G*) Effect of temperature on the onset of evening activity peaks for flies shown in *F*. One-way ANOVA with the factor of temperature. Individual differences calculated by Tukey HSD tests. Significance determined versus the 25 °C group. (*H*) Daytime, nighttime, and total activity of flies shown in *F*. Total activity (gray bar) represents group mean. Two-way ANOVA with factors of time (Light and Dark) and temperature. Individual differences calculated using the Tukey HSD test. Significance during light versus the 25 °C group. (*I*) Morning activity index versus temperature for flies shown in *F*. Morning activity index represents the proportion of activity that occurred between ZT0 and ZT5. One-way ANOVA followed by Tukey HSD test to calculate individual group differences. Error bars indicate means ±SEMs. \*\*\**P* <0.001, \**P*<0.01.

When flies were maintained at 10 °C under an identical short photoperiod, their activity profile was considerably altered (Fig. 1*A*). In addition to reducing overall activity, diapausing conditions markedly reduced the morning activity peak, and advanced the evening peak, eliminating the daytime siesta (Fig. 1*A*). To distinguish between the effects of temperature and photoperiod, we recorded the activity of flies maintained at 10 °C with a longer day. Extending the photoperiod from LD8:16 to either 12:12 or 16:8 minimally affected the activity profiles at either 25 °C or 10 °C (Fig. 1 *A–C*), demonstrating that photoperiod length does not account for the diapause activity rhythm. However, extending the photoperiod modified the timing of the evening activity peak relative to the onset of darkness. Flies maintained at 25 °C and LD8:16 initiated their evening activity peak almost simultaneously with the transition from lights on to lights off (Fig. 1 *A* and *D*; 0.8 ± 0.2 hrs before lights off), whereas under a longer photoperiod the evening activity began ∼3 hours before lights off (Fig. 1 *B-D*; 2.9 ± 0.1 hr before lights off at 12 hr light:12 hr dark; 3.2 ± 0.1 hr before lights off at 16 hr light:18 hr dark). In contrast, the evening activity peak of flies at 10 °C became more and more misaligned with the transition to lights off as the photoperiod increased (Fig. 1*D*; 4.0 ± 0.2 hr before lights off at LD8:16; 6.0 ± 0.2 hr before lights off at LD12:12; 7.4 ± 0.2 hrs before lights off at 16:8). Cool temperature eliminated the flies’ crepuscular activity rhythm—removing the morning activity peak and advancing the evening peak, such that most of their movement occurred during the midday/late afternoon (Fig. 1 *A–C*). Notably, their activity, though reduced, remained rhythmic, and the length of the photoperiod had little impact on the cool-induced conversion from crepuscular to mid-day activity. Under a short photoperiod (LD8:16;Fig. *1A*), diapausing flies’ advanced evening activity peak allowed them to fit their activity into the daytime, which could be more favorable in winter (Fig. 1*E*). These results suggest that a warm temperature primes flies for a long day and a cool temperature primes them for a short day. Furthermore, at 10 °C flies did not adapt their rhythm to a longer photoperiod.

As photoperiod does not play the predominant role in shaping the activity rhythm, we wondered whether the activity profile would change gradually with temperature, or, alternatively, whether there would be an abrupt change at diapause-inducing temperatures. To test this, we subjected flies to two additional cool temperatures (13 °C and 15 °C), as well as 20 °C and 18 °C, which are below the temperature that supports the most rapid development (25 °C), but still compatible with crepuscular activity in adult flies. For these experiments, we housed wild-type flies in LD16:8 cycles, as an extended light period helped to disambiguate morning vs. evening activity onset. As the temperature decreased, the evening activity peak shifted to earlier times (Fig. 1 *F and G*), total activity decreased (Fig. 1*H*), and the morning activity index (the proportion of daytime activity occurring from ZT0–6) diminished (Fig. 1*I*). Notably, the largest changes in all parameters occurred from 18 °C to 15 °C (Fig. 1 *F–I*). Given that 15 °C is the temperature below which egg production ceases, these results suggest that changes to circadian activity respond to the same temperatures that induce reproductive arrest. These results are consistent with recent studies that also show that reproductive arrest and recovery are more dependent on temperature than photoperiod (13, 17, 18)

### Cool temperature diminishes aversion to sleeping in the light

A striking feature of the diapause activity rhythm is the advanced timing of the evening activity peak, which initiates around midday (Fig. 1 *A*-*D*). Concomitant with this increased midday activity, is a reduction in the daytime siesta. The duration of the daytime siesta is known to be positively correlated with light intensity and temperature (23, 24), so the daytime siesta may reflect a light avoidance behavior, possibly to mitigate desiccation during hot summer days. Therefore, we wondered whether diapause-inducing temperatures might also reduce aversion to light.

To test the impact of temperature on the preference between shade versus bright light, we designed a custom behavioral arena. We housed 30 flies individually, each in a 44 mm x 6 mm enclosure with constant access to a 5% sucrose food source (Fig. 2*A*). We placed a neutral density filter along one half of each enclosure, so that flies could choose between spending time in a shaded zone (180 lux) or directly in brighter light (1700 lux) (*SI Appendix*, Fig. S1*A*). Except for the initial increase in temperature when the lights were turned on (ZT0), the temperature in the arena did not increase further during the rest of the day (ZT1-12), regardless of whether the environmental temperature was warm (*SI Appendix*, Fig. 1*B*) or cool (*SI Appendix*, Fig. 1*C*). Moreover, the temperatures of the shaded and unshaded zones were virtually identical at all times (*SI Appendix*, Fig. S1 *B* and *C*). To provide a static light source for real-time video tracking, we constantly backlit the arena with a near-IR LED light (850 nm), which is imperceptible to flies.

**Fig. 2.**
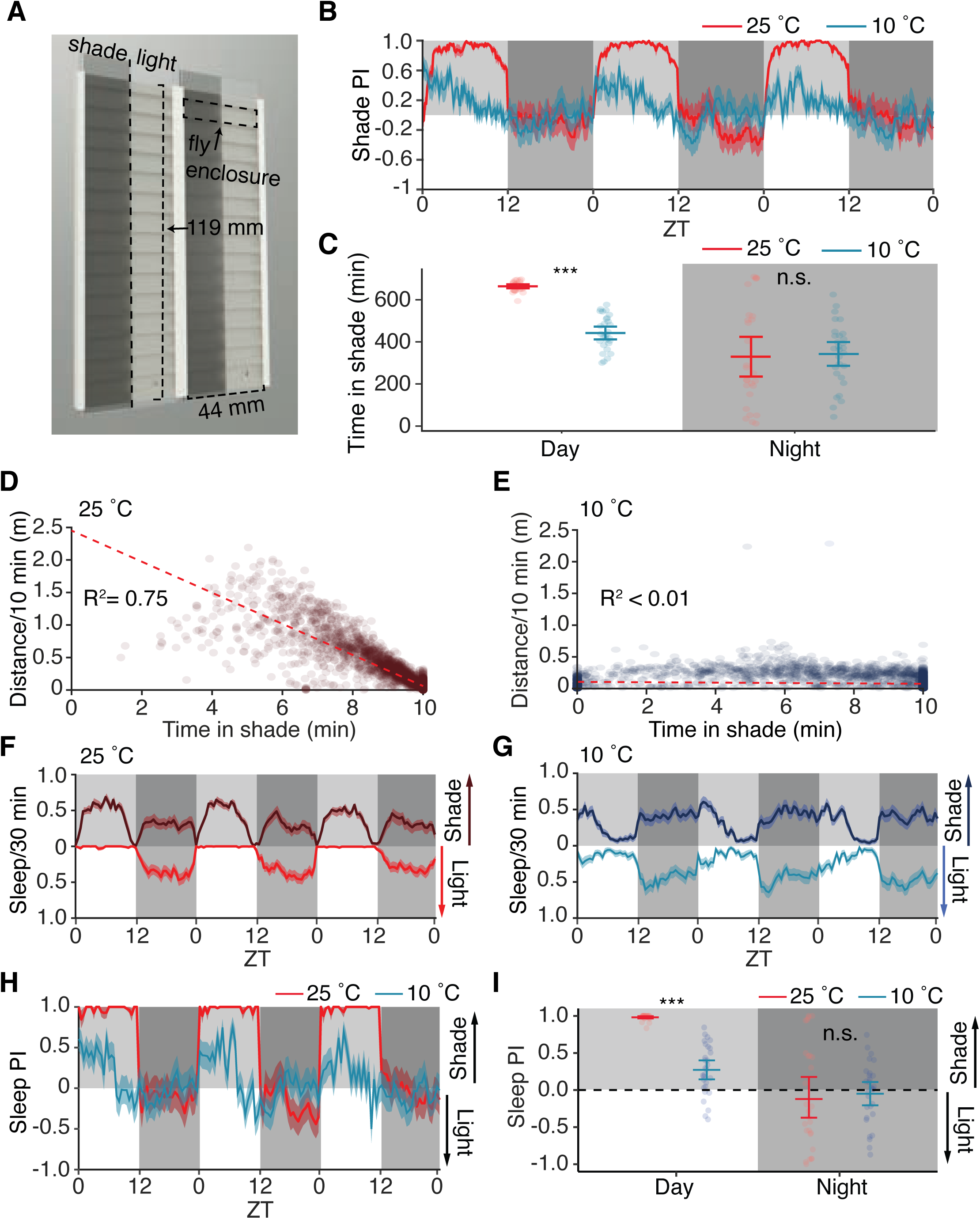
Diapause suppresses flies’ aversion to light. (*A*) Model of behavioral arena used in shade preference assay. Arena provides individually-housed flies with ad libitum access to food in each fly enclosure, as well as the choice to spend time in a shaded or unshaded zone. Fly location is tracked in real time via camera and custom software (see Methods). (*B*) Shade preference index of flies over three days at 25 °C or 10 °C. Shade preference index represents the average time each fly was in the shade or light per 10 minutes. Line represents mean shade PI, and shading around line represents SEM. (*C*) Quantification of average time spent in shade over three-day long recording during the daytime and nighttime. Error bars indicate the 95% confidence interval around the mean. Data were analyzed first by Aligned Rank Transform Two-Way ANOVA, examining factors of time, temperature, and time: temperature interaction. Individual differences between groups were analyzed by Aligned Rank Transform Contrast, with Bonferroni-adjusted *P* value. (*D*) Correlation between distance/10 minutes and time spent in the shade/10 minutes from flies at 25 °C. Each dot represents a 10 minute bin from the daytime in a three-day long recording. Red dashed line represents the linear best fit line. (*E*) Correlation between distance/10 minutes and time spent in the shade/10 minutes from flies at 10 °C. Each dot represents a 10 minute bin from the daytime in a three-day long recording. Red dashed line represents the linear best fit line. (*F*) Sleep location plot for three days from flies housed at 25 °C. Values above zero (dark red) indicate time spent sleeping in the shaded zone of the arena, while values below zero (bright red) indicate time spent asleep in un-shaded zone. Error indicates SEM. (*G*) Sleep location plot for three days from flies housed at 10 °C. Values above zero (dark blue) indicate time spent sleeping in the shaded zone of the arena, while values below zero (light blue) indicate time spent asleep in the un-shaded zone. Error indicates SEM. (*H*) Sleep preference index plot for three days from flies housed at 25 °C (red) or 10 °C (blue). Values above zero indicate a preference for the shaded zone, while values below zero indicate a preference for un-shaded zone (see Methods for formula). (*I*) Average sleep preference for flies housed in 25 °C (red) or 10 °C (blue) during daytime or nighttime. Values above zero indicate a preference for the shaded zone, and values below zero indicate a preference for the un-shaded zone. Error bars represent the 95% confidence interval around the mean. Data were analyzed first by Aligned Rank Transform Two-Way ANOVA, examining factors of time, temperature, and time:temperature interaction. Individual differences between groups were analyzed by Aligned Rank Transform Contrast with Bonferroni multiple testing correction. *B*–I: n = 27–29 Canton S (wild type flies) per condition, each housed under a 12 hr light:12 hr dark cycles. \*\*\**P*<0.001.

We found that maintaining flies at a diapause-promoting temperature (10 °C) reduced their preference for a shady environment. At 25 °C, flies ardently preferred spending time in the shaded zone during the day, especially during midday (Fig. 2 *B* and *C*). Consistent with previous reports (24, 25), we found that the flies’ preference for shade was highly correlated with the length of their immobility during the daytime (Fig. 2*D*), indicating that they were resting in the shade. In contrast to flies maintained at 25 °C, flies at 10 °C displayed a marked reduction in their preference for shade during the day (Fig. 2*B* and *C*; 664 ± 5 minutes at 25°C vs. 442 ± 15 minutes at 10 °C). Unlike flies at 25 °C, which showed a negative correlation between how far they moved vs. time spent in the shade (Fig. 2*D*), at 10 °C there was no correlation between these two parameters (Fig. 2*E*), indicating that flies at 10 °C were equally prone to rest under high or low illumination.

Because we observed that at 25 °C the flies’ preference for shade both coincided with the daytime siesta and was inversely correlated with movement, we next analyzed flies’ preference for sleeping in the shade. *Drosophila* sleep has both a daytime and nighttime component, and daytime sleep consists of shorter, lighter sleep bouts that peak around the middle of the day (26). To examine the effects of temperature on the flies’ sleep location, we used the typical definition of sleep as five consecutive minutes of inactivity (27–29). Compared to flies at 25 °C, those maintained at 10 °C exhibited a similar total amount of daytime sleep, although the timing was markedly altered such that they slept late into the morning and were more active in the afternoon (*SI Appendix*, Fig. S1 *F* and *G*). Additionally, nighttime sleep was substantially increased at 10 °C, with these flies sleeping nearly all night (*SI Appendix*, Fig. S1*F* and *G*; 424.2 ±14.1 min at 25 °C; 629.2 ± 3.9 min at 10 °C). At 25 °C, nearly all of the flies’ daytime sleep occurred in the shade (Fig. 2*F*) (30, 31). In contrast, flies at 10 °C showed only a modest preference for shaded daytime sleep during the morning, and none in the evening (Fig. 2 *G* and *H*). In total, 10 °C markedly reduced aversion to sleeping under bright illumination (Fig. 2*I*; PI =0.98 ± 0.2 at 25 °C and PI=0.27 ± 0.1 at 10 °C).

A hallmark of diapause is the arrest of oogenesis, which prevents yolk uptake, resulting in small, underdeveloped ovaries [*SI Appendix*, Fig. S2 *A* and *B*; (12)]. Ovarian arrest during diapause is in part regulated by a reduction in juvenile hormone (JH) signaling (8, 32, 33). Treating flies with the JH analog Methoprene partially reverses this arrest, causing ovaries to enlarge [*SI Appendix*, Fig. S2 *B* and *C*; (13, 34)]. To test whether the alterations we observed in flies’ circadian rhythm and sleep were also regulated by JH, we added 100 µg of the JH analog methoprene to each fly’s enclosure in our behavioral arena and recorded their activity, sleep, and preference for shade. The addition of methoprene had almost no impact on activity (*SI Appendix*, Fig S2*D*), sleep (*SI Appendix*, Fig S2*E*), or preference for shaded sleep (*SI Appendix*, Fig S2*F*). This result shows that temperature exerts JH-dependent and JH-independent effects on *Drosophila* behavior and physiology.

### Diapausing flies sleep deeply

A defining characteristic of sleep, as opposed to simple immobility, is decreased sensitivity to external stimuli (35). Because the walking speed and spontaneous movement of *Drosophila* decreases with temperature (36) (Fig. 1*F* and *H*), we wondered whether the sleep state in diapausing flies is distinct from non-diapausing flies. To address this, we compared the arousal thresholds of flies at 25 °C and at 10 °C. To perform this assay, we used vibrating motors to deliver a set of five gradually-increasing vibration stimuli, each three seconds in length, once every two hours during the course of a day (Fig. 3 *A* and *B*; SI Appendix, Fig. S3 *A* and *B*). We measured movement via real-time video tracking capable of resolving movements of less than one body-length (∼3mm)(*SI Appendix*, Video 1), and scored arousal as the vibration stimulus required to induce 3 mm of movement in flies that were immobile prior to the start of the stimulus. We observed that at 25 °C, flies readily responded to the vibration stimuli by transiently increasing their movement both during the day and at night (Fig. 3*C* and *SI Appendix*, Fig. S3*C*). Similar to flies at 25 °C, subjecting flies maintained at 10 °C to vibration stimuli during the day elicited a locomotor response (Fig. 3*D* and *SI Appendix*, Fig. S3*C*). However, the stimuli had a markedly smaller effect on the locomotion of flies at 10 °C during the night, suggesting again that these flies were in a deep sleep. Despite the vibrations transiently reducing sleep and increasing activity (Fig. 3*C* and *D*; *SI Appendix*, Fig. S3*C*), there was no overt effect of vibration stimuli on flies’ shaded sleep PI, indicating that this preference is unchanged by wake-promoting stimuli (*SI Appendix,* Fig. S3*D*).

**Fig. 3.**
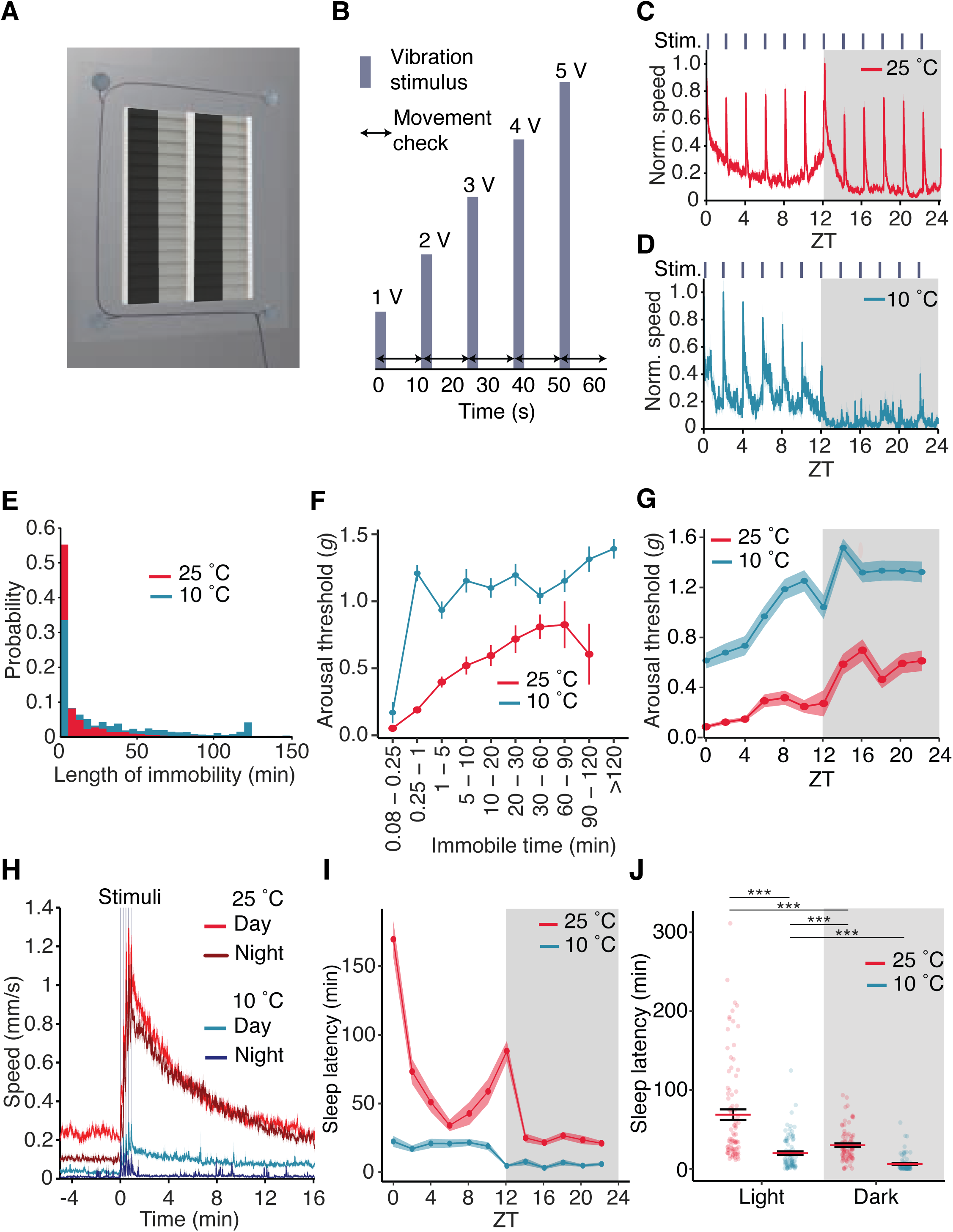
Diapause induces rapid-onset deep sleep. (*A*) Model of behavioral arena affixed with a vibrating motor at each corner. (*B*) Vibration stimulus train diagram. Every two hours, 1–5 volts were delivered to each vibrating motor, which served to increase the vibration intensity (shown in *SI Appendix,* Fig. S2*A*). Each stimulus was delivered for three seconds followed by a ten second pause. Movement was scored for arousal threshold experiments when a fly that was immobile prior to the stimulus train moved at least 3 mm, either during the vibration stimulus or in the proceeding 10 seconds. (*C*) Activity profile of flies housed at 25 °C (red) subjected to vibration stimulus train once every two hours over the span of a day. Data are binned each minute and normalized to visualize response to vibration stimulus train. (*D*) Activity profile of flies housed at 10 °C subjected to vibration stimulus train once every two hours over the span of a day. Data are binned each minute and normalized by temperature to visualize response to vibration stimulus train. (*E*) Histogram displaying probability of immobility length of flies housed in 25 °C (red) or 10 °C (blue) prior to experiencing stimulus train. (*F*) Arousal threshold vs. immobility plot. Y axis displays average arousal threshold (vibration intensity measured in *g* force) required to induce locomotion in flies at varying lengths of immobility. We did not observe at 25 °C flies that were immoble for >120 min. Error represents SEM. (*G*) Circadian arousal threshold plot displaying the mean *g* force required to induce locomotion in quiescent flies at 10 °C or 25 °C over the span of a day. E–G: Data represents 1,959 response events from 179 *w^1118^* flies. (*H*) Average walking speed of flies in response to vibration stimuli at 25°or 10°C during the day and night. Time 0 denotes the start of the stimulus train, with the vertical gray shading indicating each vibration stimulus. (*I*) Circadian sleep latency plot of flies housed at 25°C (red) or 10°C (blue). Data represent the average time to sleep (5 consecutive minutes of activity) from immobile flies that responded to the vibration stimuli throughout the day. Shading represents SEM. (*J*) Quantification of sleep latency during daytime and nighttime from flies housed at 25°C or 10°C. n=89-90 flies/temperature. Data were analyzed first by Aligned Rank Transform Two-Way ANOVA, examining factors of light, temperature (*P*<0.001), and light:temperature interaction. Individual differences between groups were analyzed by Aligned Rank Transform Contrast with a Bonferroni multiple testing comparison. \*\*\**P*<0.001.

Sleep depth (arousal threshold) in *Drosophila* changes with the length of their immobility, suggesting that, as in mammals, fly sleep may also have unique stages, and therefore have sleep architecture (i.e, distinct periods of sleep characterized by different arousal thresholds) (35, 37, 38). To assess whether diapause impacts sleep architecture, we analyzed how their arousal threshold changed with their length of immobility. We found that flies maintained at 10 °C were, on average, immobile for ∼10-fold longer than flies at 25 °C prior to the onset of the vibration stimuli (Fig. 3*E*; median time of 1.5 ± 0.6 minutes at 25 °C vs. 14.3 ± 1.1 minutes at 10 °C). At 25 °C, the arousal threshold, which is the vibration intensity (expressed in units of gravitational force, *g*), increased greatly if the flies were immobile for a long time. After 5–15 seconds of immobility, non-diapausing flies could be aroused easily with low vibration intensity (Fig. 3*F*; 0.05 ± 0.01 *g*). After 5–10 minutes, 10-fold more *g* force was required to stir the flies (Fig. 3*F*; 0.52 ± 0.06 *g*). The force required peaked at 60–90 minutes of immobility (Fig. 3*F*; 0.83 ± 0.18 *g*), after which it decreased (Fig. 3*F* and *SI Appendix*, Fig. S3*E*; 0.61 ± 0.22 *g*).

We found that the arousal threshold of flies at 10 °C exhibited a different pattern. 10 °C did not simply impair their ability to sense vibration because flies that were immobile for 5–15 seconds had a low arousal threshold similar to flies at 25 °C (Fig. 3*F*; 0.17 *g*). However, at immobilities greater than 15 seconds, the arousal threshold of diapausing flies rapidly increased. After only 15–60 seconds of immobility, the arousal threshold of diapausing flies (1.21 ± 0.6 *g*) exceeded even the greatest value that we observed in flies at 25 °C (Fig. 3*F*). Also unlike flies at 25 °C, the arousal threshold of diapausing flies did not peak after 60-90 minutes of immobility, but remained elevated, even in flies that had been immobile for >2 hours (Fig. 3*F*). These results indicate that diapausing flies rapidly initiate a long-lasting, deep sleep state that is distinct from non-diapause sleep.

### Flies awakened in diapause rapidly resume sleep

To assess the circadian influence on sleep depth, we analyzed how the arousal threshold changed throughout the day at 25 °C and 10 °C. During the day, flies at 25 °C had their highest arousal threshold (i.e. deepest sleep) during the period of their midday siesta (Fig. 3*G*), and an overall higher arousal threshold at night (Fig. 3*G*). In contrast, the daytime arousal threshold of flies at 10 °C started out 6-fold higher and further increased during the day, with their deepest sleep occurring at ∼ZT 10 (Fig. 3*G*). Unlike flies at 25 °C, the proportion of diapausing flies responding to vibration stimuli displayed a nearly monotonic decrease as the day progressed (*SI Appendix*, Fig. S3*F*), suggesting that a cool temperature rapidly increased sleep pressure throughout the day. However, similar to flies at 25 °C, flies at 10 °C had a higher arousal threshold and lower proportion of responders at night compared to the day (Fig. 3*G* and *SI Appendix*, Fig. S3*F*). Overall, flies at 10 °C exhibited rhythmic circadian sleep patterns, but they were profoundly different from those of flies at 25 °C.

We reasoned that if a cool temperature increased sleep pressure, then there would be differences in how quickly diapausing and non-diapausing flies resume sleep after having been startled by the vibration stimulus. To address this, we plotted how walking speed changed in response to vibration. The baseline walking speed, which is the average walking speed prior to a vibration stimulus, was lower during the night than during the day at both 25 °C and 10 °C (Fig. 3*H* and *SI Appendix*, Fig. S3*G*). Upon vibration, flies at 25 °C, on average, rapidly increased their walking speed from baseline (0.23 mm/s day; 0.10 mm/sec night) to a maximum of 1.10 mm/sec during the night and 1.29 mm/sec during the day (Fig. 3*H* and *SI Appendix*, Fig. S3*G*). After the stimuli, their average daytime and nighttime walking speed gradually decayed back to the baseline at roughly the same rate over 16 minutes (*SI Appendix*, Fig. S3*G*). Flies at 10 °C also increased their walking speed in response to the vibration stimuli; however, during the nighttime, diapausing flies rapidly stopped moving (Fig. 3*H* and *SI Appendix* Fig. S3*G*).

To quantify this effect further, we recorded sleep latency for diapausing and non diapausing conditions. We defined sleep latency as the time it takes for immobile flies that are responsive to the vibration stimuli to initiate an extended bout of rest (5 consecutive minutes of immobility). The sleep latency of flies at 25 °C dramatically changed throughout the day, with their longest sleep latency occurring in the morning and early night (Fig. 3*I*; 169 ± 13 min and 88 ± 7 min, for ZT 0 and ZT 12.1, respectively), and their shortest daytime latency occurring at midday (ZT 6; 34 ± 4 min) (Fig. 3*I*). Late in the night (ZT 14–22), the sleep latency of flies at 25°C was more consistent, averaging ∼25 minutes. In contrast, the sleep latency of diapausing flies was markedly reduced, both during the daytime (average for ZT 0–10: 69 ±7 min at 25 °C vs. 20 ± 2 min at 10 °C) and nighttime (average for ZT 12.1–22: 30 ±2 min at 25 °C vs. 6 ±1 min at 10 °C). The sleep latency of flies at 10°C fluctuated little over the course of the day or night (Fig. 3 *I* and *J*); however, it was still higher during the day compared to the night, showing again that the flies maintained rhythmic behavior.

These data demonstrate that flies in diapause rapidly resume sleep upon being startled, and highlight the soporific effects of cool temperature on flies. In total, the arousal threshold data reveal that diapausing flies enter a unique, deep-sleep state, characterized by a rapid onset, high arousal threshold, altered cycling characteristics, and rapid resumption of sleep following awakening.

### The eye and cryptochrome are dispensable for shaded sleep preference

We next sought to understand the genetic basis for the shaded sleep preference at 25 °C. We reasoned that two classes of genes could be important for this behavior: those that comprise the circadian clock, since the clock determines the timing of the daily siesta, and those that are involved in light sensation. To test the latter, we first examined the shaded sleep preference of flies with major defects in visual transduction in the compound eye.

The compound eye is composed of an array of ∼800 hexagonal ommatidia, each of which houses eight photoreceptor cells. The phototransduction cascade is initiated by light activation of rhodopsin, engagement of a heterotrimeric G-protein, and subsequent activation of a phospholipase Cb encoded by the *norpA* locus (39). The cascade then culminates with opening of two cation channels, TRP and TRPL, causing depolarization of the photoreceptor cells (40–42). Surprisingly, flies harboring null mutations in *norpA* or double mutant for *trp* and *trpl* (*trpl^MB10553^*;*trp^MB03672^*) showed a strong daytime preference for shaded sleep at 25 °C that was not significantly different from the wild-type control (Fig. 4 *A* and *B*; control PI=0.84 ±0.04, *trpl^MB10553^*;*trp^MB03672^* PI=0.82 ±0.04, *norpA^P24^* PI=0.96±0.04).

**Fig. 4.**
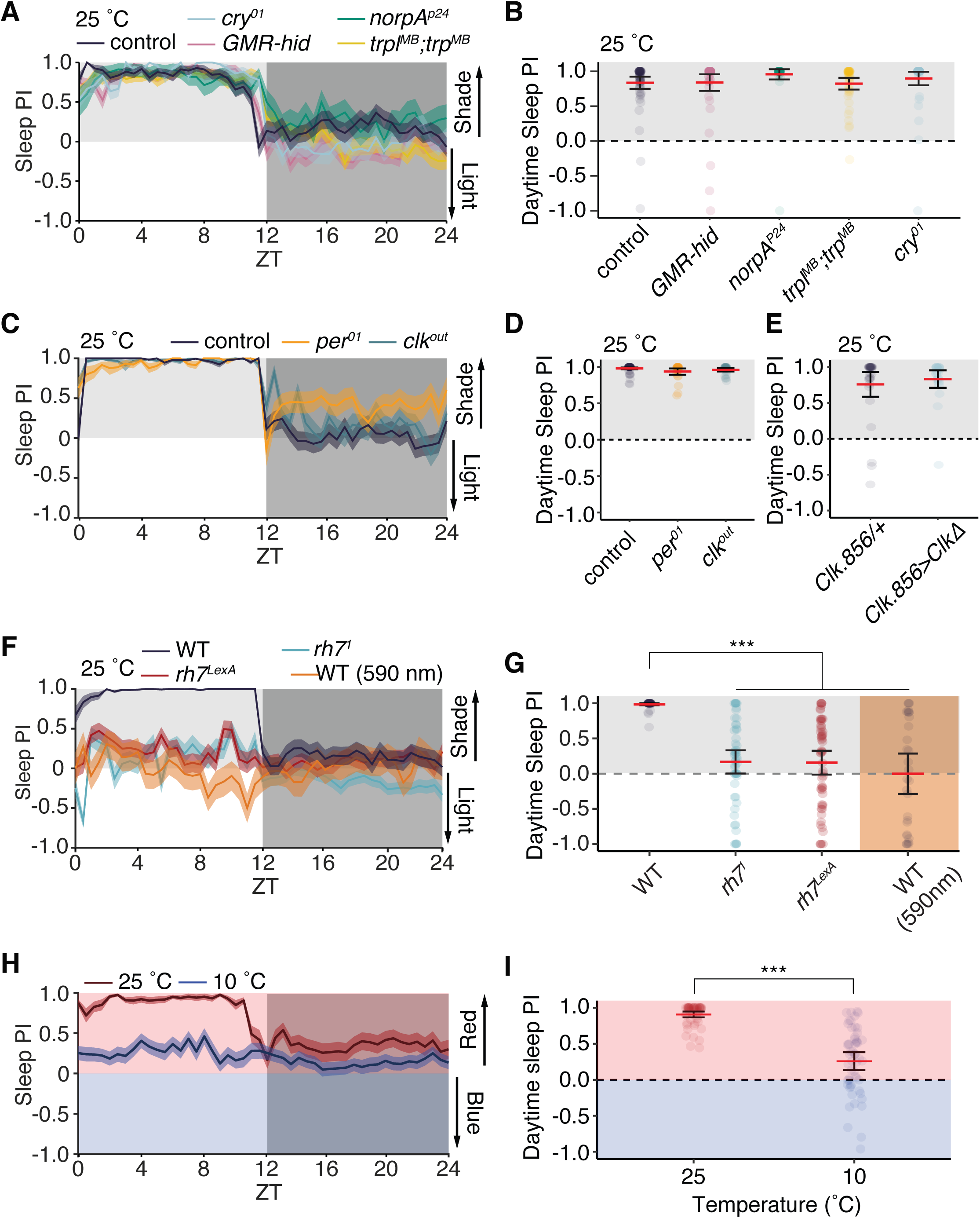
Rh7 is required for flies’ shaded sleep preference. (*A*) Sleep preference index plot for flies with major defects in the visual system or Cryptochrome at 25 °C. Values above zero indicate a preference for the shaded zone, while values below zero indicate a preference for un-shaded zone (see Methods for formula). (*B*) Average sleep preference for flies shown in panel A during the daytime. Values above zero indicate a preference for the shaded zone, and values below zero indicate a preference for the un-shaded zone. Error bars represent the SEM. Data were analyzed first by Aligned Rank Transform Two-Way ANOVA. Individual differences between groups were analyzed by Aligned Rank Transform Contrast. *A* and *B*: n=45–59 flies/genotype. (*C*) Sleep preference index plot for flies with disruptions to the endogenous circadian clock at 25°C. Values above zero indicate a preference for the shaded zone, while values below zero indicate a preference for un-shaded zone (see Methods for formula). (*D*) Average sleep preference for flies shown in panel C during the daytime. Values above zero indicate a preference for the shaded zone, and values below zero indicate a preference for the un-shaded zone. Error bars represent SEM. Data were analyzed first by Aligned Rank Transform Two-Way ANOVA. Individual differences between groups were analyzed by Aligned Rank Transform Contrast. *C* and *D*: n=27–58 flies/genotype. (*E*) Average sleep preference for flies expressing dominant negative CLK (CLKΔ) in circadian pace-maker neurons. n=24–29 flies/genotype. (*F*) Sleep preference index plot for flies with mutations in *rh7* or wild-type flies housed under red-shifted (590 nm) light. Values above zero indicate a preference for the shaded zone, while values below zero indicate a preference for un-shaded zone (see Methods for formula). (*G*) Average sleep preference for flies shown in panel F during the daytime. Values above zero indicate a preference for the shaded zone, and values below zero indicate a preference for the un-shaded zone. Error bars represent SEM. Data were analyzed first by Aligned Rank Transform Two-Way ANOVA (*P*<0.001). Individual differences between groups were analyzed by Aligned Rank Transform Contrast. *F* and *G*: n=30–56 flies/genotype. (*H*) Red vs. blue sleep preference index plot for flies housed at 25 ° or 10 °C. (*I*) Quantification of daytime sleep preference index from panel H. Data were analyzed first by Aligned Rank Transform One-Way ANOVA. n=54 flies/temperature. *P*<0.001.

In addition to the two compound eyes, flies possess three small eyes at the vertex of the head (ocelli) (43), as well as two Hofbauer–Buchner (H-B) eyelets, which are located between the retina and optic lobes (44). To test whether any eye structure is important for shaded sleep preference, we took advantage of flies that express a pro-apoptotic gene (*head involution defective*, *hid*) under the control of the *GMR* (*Glass Multimer Reporter*) promoter (45, 46), thereby resulting in elimination of the compound eyes, ocelli and H-B eyelets. We examined the shaded sleep preference of *GMR-hid* flies, and found that they retained a strong preference for daytime shaded sleep (Fig. 4*A* and *B*; PI=0.84 ±0.06). These results demonstrate that eye structures are not required for light aversion.

In addition to sensing light through eye structures, *Drosophila* can also detect light outside of the eye via the flavoprotein, Cryptochrome (Cry) (47–49). Cry senses ultraviolet/blue light (450 nm peak) (50) and plays a role in phase-shifting the endogenous molecular clock in response to light, as well as the avoidance of UV light. However, flies with a null mutation in *cry* retained a strong preference for shaded sleep (Fig. 4 *A* and *B*; PI=0.9 ±0.04).

### A functional circadian clock is not required for shaded sleep preference

To test whether the circadian clock impacts shaded sleep preference, we examined shade preference in clock mutants. The *Drosophila* circadian clock consists of at least two interlocked transcription-translation negative feedback loops. The positive limb of the primary clock loop consists of the transcription factors CLOCK (CLK) and CYCLE (CYC), which heterodimerize to regulate the transcription of *period* (*per*) and *timeless* (*tim*) (51). In turn, PERIOD and TIMELESS heterodimerize, translocate into the nucleus and repress transcription of the *clock* and *cycle* genes. Movement through this cycle takes ∼24 hours, and disrupting any of the transcription factors mentioned above renders flies behaviorally arrhythmic when housed in conditions devoid of time-giving cues, such as light.

To test whether the molecular clock regulates the preference for shaded sleep, we tested flies with null mutations in either *per* (*per^01^*) or *Clk* (*Clk^OUT^*). Similar to the control, both mutants showed a high preference for sleeping in the shade (Fig. 4*C* and *D*; control PI=0.98 ±0.01; *per^01^* PI=0.94 ±0.02; *Clk^OUT^* PI=0.96 ±0.01). Similarly, disrupting the molecular clock in the pacemaker neurons (the neurons in the brain that drive 24-hour activity rhythms) by expressing a dominant negative CLOCK protein (52) failed to suppress the preference for sleeping in the shade (Fig. 4*E*; *clk.856>ClkΔ* PI=0.83 ±0.1; *clk.856-Gal4*/+ PI=0.76 ±0.1). Together, these results demonstrate that a functional circadian clock is not required for the flies to prefer shaded sleep.

### Rhodopsin 7 regulates preference for shaded sleep

In addition to CRY, another extraocular light sensor is encoded by the *rhodopsin 7* (*rh7*) gene, which is expressed in the brain (53) and in multidendritic neurons (54). This blue-light-sensitive opsin impacts circadian entrainment to blue light, but does not signal through the phospholipase Cβ encoded by the *norpA* locus (53). Because we observed that *norpA* mutants retained a strong preference for shaded sleep, we next tested whether Rh7 regulates this behavior. We tested the sleep preference of flies homozygous for either of two null alleles–*rh7^1^* (*^53^*) and a new allele, *rh7^LexA^*, which contains *LexA* and *mini-white* positioned in frame following the endogenous initiation codon in the second exon of *rh7* (*SI Appendix*, Fig. S3*H*). Compared to the wild-type control, which showed a strong preference for shaded sleep (Fig. 4 *F* and *G*; PI=0.99 ±0.01), both *rh7* mutants showed only a minimal preference for sleeping in the shade at 25 °C (Fig. 4 *F* and *G*).

Rh7 is activated by blue light (53), so we next tested the sleep of flies housed in amber light (λ=591 nm), which is outside of the spectral sensitivity of Rh7. Unlike flies housed under a broad-spectrum white light, flies under the amber light showed no preference for sleeping in the shade (Fig. 4*G*; PI=0 ±0.14). Together, these results demonstrate that the preference for shaded sleep is specific for blue-light, is independent of all eye structures and the circadian clock, and requires Rh7.

### Cool temperature suppresses *rh7-*mediated aversion to blue light

To directly test whether shaded sleep preference is driven by avoidance of blue light, we modified our behavioral arena by placing a blue filter over one half of the arena and a red filter over the other, such that flies could choose to sleep in a red or blue zone (*SI Appendix*, Fig. S4*A*). The red filter allowed peak emission of λ=612 nm, with virtually no emission below λ=575 nm (*SI Appendix*, Fig. S4*B*). The peak emission with the blue filter was 448 nm, although the spectrum had a smaller peak at λ=510 nm, and narrow peak at λ=612 nm (*SI Appendix*, Fig. S4*B*). Flies maintained at 25 °C strongly preferred sleeping in the red-shaded half of the arena (Fig. 4 *H* and *I*; PI=0.91 ±0.02), reminiscent of flies choosing to sleep in the shade rather than in white light. However, at 10 °C flies showed only a minimal aversion to sleeping under blue light (Fig. 4 *H* and *I*; PI=0.26 ±0.06). This suggests that the indifference to shaded sleep in the cold is the result of insensitivity to blue light.

To test whether the activation of *rh7* positive neurons drives aversion to blue light, we used *rh7^LexA^* to express a transgene encoding a red-light-sensitive channelrhodopsin, CsChrimson (55), in *rh7*-positive neurons. We housed the flies in the same arena where they could choose between spending time in a red-or blue-shaded zone. However, in this paradigm, we activated *rh7*-positive neurons on both halves of the arena. In the red-shaded zone they were activated by CsChrimson, and in the blue-shaded zone by endogenous Rh7. Because CsChrimson requires the cofactor all-*trans* retinal (ATR) to be activated by red light, we compared the light preference of flies that had been raised on food with or without 1 mM ATR. Control flies (i.e., *rh7^LexA^*>*CsChrimson* flies raised on food without retinal) avoided spending time in blue light throughout the day (Fig. 5*A*), with only 215 ±15 minutes, out of a possible 720 minutes, spent outside of the red zone (Fig. 5*B*). In contrast, flies with functional CsChrimson avoided the red half of their enclosure, more than doubling the amount of time spent under blue light (492 ±16 minutes; Fig.5*B*). Strikingly, we observed a similar trend for sleep preference. The control without ATR strongly preferred sleeping on the red side of their enclosure (PI= –0.83 ±0.06; Fig. 5*C*). In contrast, activating *rh7*-positive neurons with CsChrimson nearly reversed this preference, as these flies preferred to sleep under blue light (PI=0.42 ±0.1; Fig. 5*C*). In total, these results reveal that the activation of *rh7*-positive neurons is sufficient to change the valence of light with respect to sleep, as Rh7 activity drives an aversive response.

**Fig. 5.**
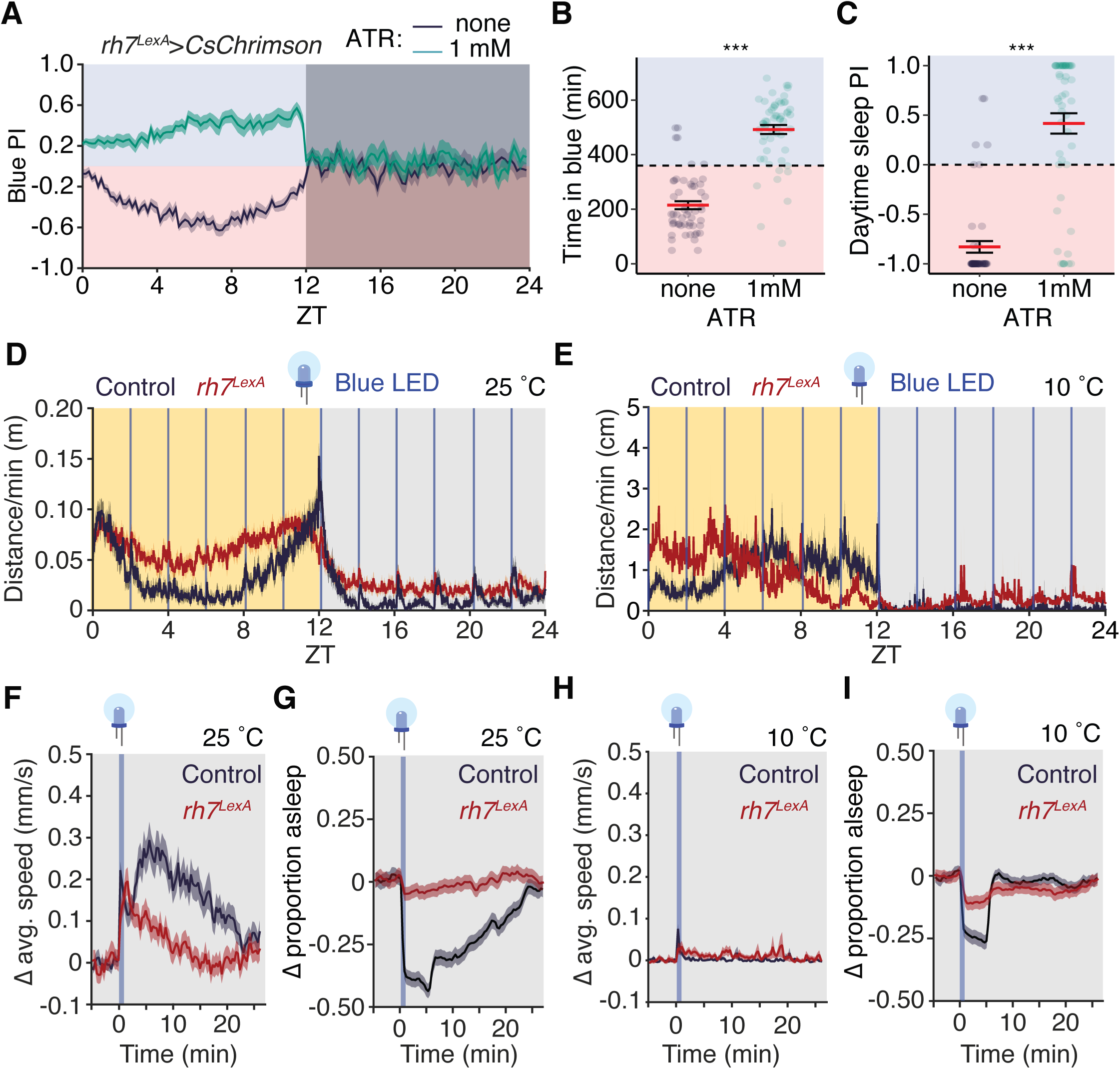
A cool temperature suppresses the wake-promoting effects of blue light. (*A*) Blue preference index plot for *rh7>*csChrimson flies raised on regular fly food not supplemented with all-trans retinal (none) or food supplemented with 1 mM all-trans retinal (1 mM). Shading indicates SEM. (*B*) Total time spent in the blue portion of enclosure during the day for flies shown in panel A. Errorbars indicate SEM. (*C*) Average daytime preference index for sleeping in the blue portion of enclosure for flies shown in panel A. Errorbars indicate SEM. B and C Data were analyzed using one-way Aligned Rank Transform ANOVA. n=54–57 flies per condition. (*A*–*C*.) Activity profile of control and *rh7^LexA^* flies exposed to blue-light pulses once every two hours over the span of a day. (*D*) Flies maintained at 25 °C. (*E*) Flies maintained at 10 °C. (*F*) Change in average speed of flies exposed to blue light pulses during the night when maintained at 25 °C. Data normalized to pre-stimulus baseline. (*G*) Change in proportion of flies asleep after blue light pulses given at night when maintained at 25 °C. Data normalized to pre-stimulus baseline. (*H*) Change in average speed of flies exposed to blue light pulses during the night when maintained at 10 °C. Data normalized to pre-stimulus baseline. (*I*) Change in proportion of flies asleep after blue light pulses given at night when maintained at 10 °C. Data normalized to pre-stimulus baseline. F–I represent the average of each late-night blue light pulse (ZT 14–22). Error indicates SEM. n=50–59 flies/condition. \*\*\**P*<0.001.

### Cool temperature suppresses the wake-promoting effect of blue light

Non-diapausing flies might avoid sleeping in blue light in part because blue light promotes wakefulness, and flies are unable to fall asleep when exposed to it. To test this hypothesis, we housed flies in 12 hr light:12 hr dark cycles, with daytime light from a 591 nm light source, which is outside of the spectral sensitivity of Rh7 {Ni, 2017 #5598}. Every two hours, we delivered five three-second pulses of blue light and measured the behavioral responsiveness to these stimuli.

At 25 °C, control flies responded to nighttime blue-light pulses with a stereotypical movement pattern. These flies exhibited an immediate startle response to blue light (i.e., an increase in activity that was coincident with the start of the light pulses) followed by an elevation in activity that took ∼25 minutes to return to baseline (Fig. 5 *D* and *F*). There was a commensurate reduction in sleep in these flies with, on average, a 45% reduction in sleep immediately following the light pulse (Fig. 5*G*). The *rh7* mutant flies also exhibited a startle response to blue light; however, their activity levels rapidly returned to baseline (Fig. 5*F*), suggesting that *rh7* activity is required for the prolonged stimulatory effects of blue light. Blue light pulses failed to change the proportion of *rh7* mutant flies that were asleep, suggesting that Rh7 activity underlies blue light’s wake-promoting effects (Fig 5*G*).

Unlike at 25 °C, control flies at 10 °C were relatively indifferent to nighttime blue light pulses (Fig. 5*E*). Although there was a minor startle effect from blue light, as evidenced by the small, transient increase in their average walking speed, the movement of these flies rapidly returned to baseline (Fig. 5*E* and *H*). Similarly, blue light did not substantially reduce sleep in flies maintained at 10°C, with a rapid (<10 minutes) resumption of sleep in flies that were awoken by the light stimulus (Fig. 5*I*). The change in activity and sleep in *rh7* mutant flies subjected to blue light pulses at 10°C was nearly indistinguishable from the wild type control (Fig. 5 *H* and *I*). In total, these results reveal that a cool temperature can overcome the stimulatory and wake-promoting effects of blue light, which likely contributes to the relative indifference to sleeping in the shade at 10 °C.

### Diapause conditions induce neuronal markers of high sleep pressure

Like in mammals, sleep in *Drosophila* can be described by a two-process model consisting of the internal circadian clock, which affects the timing of sleep throughout the day, and the sleep homeostat, which imparts sleep pressure as a consequence of the previous length of wakefulness (56). Notably, flies experiencing high levels of sleep pressure (such as those that have been sleep deprived for a night) will respond by increasing their total levels of sleep, and by experiencing deeper sleep than normal (57, 58). Because we observed that flies at a diapause-permissive temperature exhibit behavioral characteristics of a state of high sleep pressure (need) including an increase in arousal threshold and total sleep, as well as a decrease in sleep latency, we wondered whether they display increased expression of neuronal markers associated with sleep drive.

To assess expression of sleep pressure markers, we first examined Bruchpilot (BRP), a presynaptic active-zone protein that drives sleep need (pressure) and sleep depth (59). We housed flies at either 25 °C or 10 °C for ∼36 hours and at ZT 4 stained dissected brains with anti-BRP. In wild-type flies maintained at 10 °C, BRP expression was elevated two-fold relative to flies at 25 °C (Fig. 6 *A* and *B*; 7.5 ±1.2 AU at 25 °C; 15.4 ±1.4 AU at 10 °C). Similarly, BRP expression was also increased in *rh7* mutant flies in response to cold (*SI Appendix*, Fig. S4 *C*–*E*; 2.9 ±0.3 AU at 25 °C; 9.9 ±1.2 AU at 10 °C;). This demonstrates that the deep-sleep-like behavior observed in diapause conditions is also accompanied by a neuronal driver of high sleep pressure.

**Fig. 6.**
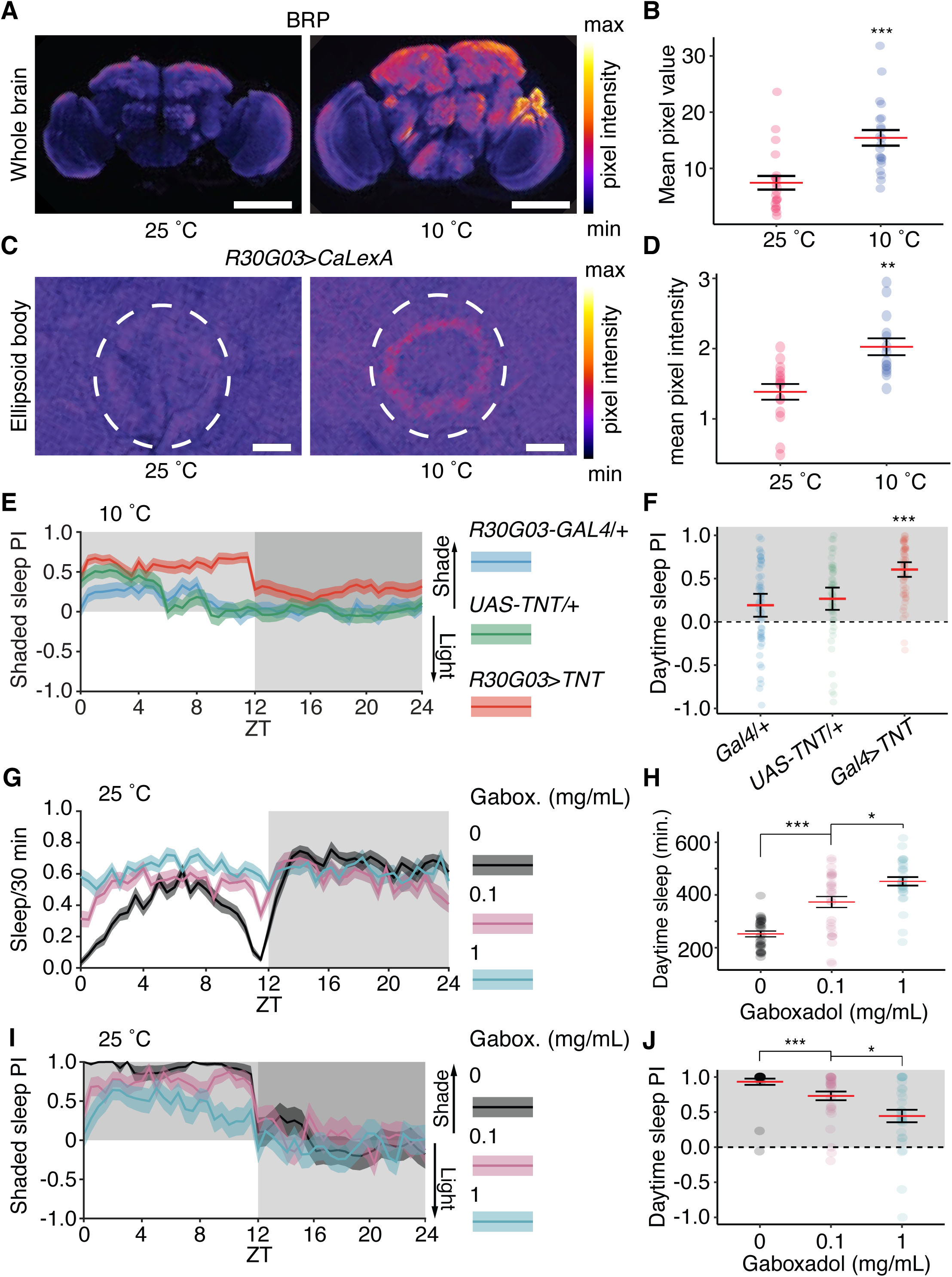
Diapause actively imparts sleep pressure, which is sufficient to overcome aversion to light. (*A*) Representative images displaying BRP staining in wildtype flies maintained at 25°C or 10 °C. Scale bar indicates 100 µm. (*B*) Quantification of BRP staining shown in A. Data were compared via Wilcoxon rank sum test. n=21 brains/temperature. (*C*) Representative images displaying R30G03-Gal4/+>CaLexA/+ from flies maintained at 25 °C or 10 °C. Scale bar indicates 10 µm. (*D*) Quantification of pixel intensity from *R30G03-Gal4*/+>*CaLexA*/+ flies maintained at 25 °C or 10 °C. Data analyzed via Wilcoxon rank sum test. n=14–15 brains/temperature. (*E*) Sleep preference index of *R30G03*>*TNT* flies at 10 °C. (*F*) Quantification of daytime sleep preference index of *R30G03*>*TNT* flies at 10 °C. Data analyzed by Aligned-Rank Transform ANOVA with group differences compared via Aligned Rank Transform Contrasts with a Bonferroni multiple testing correction. *E* and *F*: n=52–56 flies/genotype. (*G*) Sleep profile of wild-type flies fed 0, 0.1, or 1 mg/mL of gaboxadol (Gabox.) (*H*) Total daytime sleep of wild-type flies given gaboxadol. Errorbars indicate SEM. Data compared via Aligned-Rank Transform ANOVA with group differences compared via Aligned Rank Transform Contrasts with a Bonferroni multiple testing correction. (*I*) Sleep preference index of wild-type flies given gaboxadol. (*J*) Quantification of daytime sleep preference index of wild-type flies given gaboxadol. Data compared via Aligned-Rank Transform ANOVA with group differences compared via Aligned Rank Transform Contrasts with Bonferroni multiple testing correction. *G*–*J*: n=29–30 flies/genotype. \*\*\**P*<0.001.

Furthermore, these results show that *rh7* mutants accumulate sleep pressure in a manner that is similar to wild-type flies in diapause conditions.

Because BRP is reported to drive sleep pressure in part by activating R2 neurons in the ellipsoid body (60), we wondered whether flies in diapausing conditions showed elevated R2 neuron activity. To address this question, we used a genetically-encoded Ca^2+^ reporter, CaLexA, which consists of the LexA transcription factor fused to the Ca^2+^-responsive element NFAT (61). Sustained elevation of Ca^2+^ causes LexA::NFAT to translocate into the nucleus and drive expression of *LexAop-*GFP, allowing GFP expression to serve as an indirect marker of neuronal activity. We observed that flies maintained in diapausing conditions for ∼36 hours showed elevated levels of GFP in R2 cells, suggesting that 10 °C increases the activity of these neurons (Fig. 6 *C* and *D*). Notably, the activity of R2 neurons within the ellipsoid body plays a crucial role in generating sleep drive within the fly (62), indicating that the state is actively maintained rather than a passive effect of cool temperature and providing a potential mechanistic explanation for the diapause deep-sleep state.

### Sleep pressure is sufficient to overcome flies’ aversion to light

Because flies in diapausing conditions exhibit both behavioral and neuronal markers of high sleep pressure, we wondered whether an elevated level of sleep pressure was sufficient to suppress the preference for shaded sleep. To address this idea, we first expressed tetanus toxin (*UAS-TNT*) under the control of the R2 neuron driver *R30G03*-*Gal4*, which inhibits neurotransmitter release in these cells and is reported to partially alleviate sleep pressure (62). At 10 °C, *R30G03*>*TNT* flies slept significantly less per day compared to the controls (*SI Appendix,* Fig. S4 *F* and *G*; *R30G03-Gal4*/+, 870.8 ±18.2 min; *UAS-TNT*/+, 827.4 ±19.4 min; *R30G03*>*TNT*, 709.4 ±19.4 min), and had an elevated preference for shaded daytime sleep (Fig. 6 *E* and *F*; *R30G03-Gal4*/+, PI=0.19 ±0.07; *UAS-TNT*, PI=0.27 ±0.06; *R30G03*>*TNT*, PI=0.60 ±0.04).

To determine whether increasing sleep pressure through a mechanism independent of low temperature would overcome the aversion to sleeping in the light, we added a GABA_A_ receptor agonist, gaboxadol, to the food of flies maintained at 25 °C (63). Expectedly, the gaboxadol increased total sleep in a dose-dependent fashion, especially during the day, with flies fed 0.1 mg/mL of gaboxadol sleeping for 373 ± 21 minutes during the day and flies fed 1 mg/mL sleeping for 452 ± 16 minutes, versus 252 ± 11 minutes in flies fed no gaboxadol (Fig. 6 *G* and *H*). Accompanying this increase in total sleep, was a decrease in their preference for shaded sleep. Wild-type flies maintained on food without gaboxadol displayed a strong preference for shaded sleep (Fig. 6 *I* and *J*; PI=0.93 ±0.04). Whereas flies fed 0.1 mg/mL of gaboxadol had a PI of 0.73 ±0.06, with a further reduction to a PI=0.44 ±0.09 in flies fed 1 mg/mL (Fig. 6 *I* and *J*). Together, these results demonstrate that the preference for shaded sleep can be modulated by sleep drive since alleviating sleep pressure by blocking R2 neurons increased shaded sleep at 10 °C, and enhancing sleep pressure with gaboxadol dissipated this preference at 25 °C.

Notably, gaboxadol treatment did not reshape the sleep pattern such that it mimicked flies at 10 °C (Fig. 6*G* vs. *SI Appendix*, Fig. S1*F* 10 °C). Gaboxadol-treated flies exhibited slightly increased sleep during the middle of the day, whereas flies in diapause were most active during this time (Fig. 1*D*). This indicates that the diapause activity/sleep rhythm is not solely based on increased sleep pressure, further highlighting the multifarious effects of cool temperature.

## Discussion

Here we report that the same range of low temperatures (10-15 °C) that is known to induce reproductive arrest and extend lifespan in various *Drosophila* species has a strong effect on circadian behavior and sleep, independent of juvenile hormone. Cool temperatures do not trivially reduce all neuronal activity, or global levels of transcription, or simply immobilize the animals. Rather, 10-15 °C actively increases sleep pressure, causing flies to rapidly fall into a deep sleep from which it is difficult to rouse them. Nevertheless, they maintain rhythmic behavior under LD cycles, though dramatically altered from non-diapausing flies. Their sleep preferences change too–they no longer prefer sleeping in the shade over a well-lit environment. Furthermore their sleep is so deep that their arousal threshold does not seem to decrease no matter how long they have been asleep. These features are somewhat reminiscent of hibernating mammals, which show extended bouts of non-rem sleep with intermittent periods of wakefulness (3).

### Basis for the profoundly altered daytime activity pattern of diapausing flies

One of the major and unexpected discoveries uncovered by this work is that the circadian activity pattern of diapausing flies is strikingly different from non-diapausing flies. It is well established that non-diapausing flies have peak activities near dawn and dusk, and this crepuscular behavior is under circadian control (21, 51). However, we found that the peak daytime activity pattern of diapausing flies is opposite to non-diapausing flies. In contrast to non-diapausing flies that undergo a lull in activity in the middle of the day known as the siesta (51), flies maintained at 10 °C were most active during the midday. Nevertheless, the total daytime activities exhibited by diapausing and non-diapausing flies are similar. In contrast to their daytime pattern, diapausing flies show a profound reduction in their nighttime activity, relative to non-diapausing flies. Thus, diapause does not simply cause an overall reduction in movement, but profoundly shifts the circadian activity patterns.

We propose that the midday activity peak of flies at 10 °C reflects a change in circadian activity induced by the diapause program rather than an acute response to temperature. The activity pattern of the diapausing flies does not require actual changes in temperature during the day since we maintained the flies at a constant 10 °C temperature throughout the light/dark cycles. Furthermore, we found that there was an abrupt transition towards a single activity peak when we maintained the flies at ≤15 °C, which induces reproductive diapause, compared to 18 °C, which does not.

The midday activity in diapausing flies appears to be due to advancement of the evening activity peak, accompanied by an elimination of the morning activity. Consistent with this proposal, the single activity peak is not precisely in the middle of the day but shifted slightly towards dusk. This was most apparent when we maintained diapausing flies under 16 hour day and 8 hour night cycles. Under these extended day cycles, there was partial overlap of the activity peak with the evening peak for non-diapausing flies, but no overlap with the morning peak.

In the case of non-diapausing flies, the siesta is thought to reduce the risk of dehydration during the hottest part of the day. However, cool temperatures reduce that risk, offering a plausible physiological explanation and selective advantage for increased activity during the middle of the day.

### Sleeping in the shade depends on an extraocular roles for Rhodopin 7

We uncovered another difference in daytime behavior between diapausing and non-diapausing flies that we suggest might also minimize dehydration in flies at 25 °C. Using a newly developed assay, we found that non-diapausing flies have a strong preference to sleep in a shady environment, and this bias is dramatically reduced in diapausing flies. Moreover, to avoid light during daytime sleep, we found that non-diapausing flies employ an unconventional, eye-independent light-detection mechanism that depends on Rh7. In support of this model, non-diapausing flies that are missing all eye structures still seek out a shady environment for daytime sleep. This raised the question as to the identity of the extraocular light receptor that enables non-diapausing flies to avoid light and rest in a shady environment. Cry is a well-known extraocular light receptor, which is expressed in central pacemakers in the brain and functions in photoentrainment of circadian rhythms (48, 49, 64). However, the *cry* mutant was just as effective as control flies in selecting the shady zone for sleep. Thus, neither an eye structure nor Cry is required for shade preference. Rather, we found that Rh7, a light receptor that we previously showed is expressed in the brain {Ni, 2017 #5598}, is required for choosing the shade over the brightly lit area for sleep.

There are at least two non-mutually-exclusive possibilities that could explain why flies have such a substantial preference to sleep in the shade. The first is that sleeping in a lighted environment is aversive to non-diapausing flies, and if given the option, flies will avoid spending substantial amounts of time in it. Rh7 is maximally sensitive to blue light, and when given choice between zones with either red or blue light, non-diapausing flies avoid sleeping under blue lights. The second possibility is that light promotes wakefulness, and flies are unable to fall asleep when exposed to it. We found that when we use red light to stimulate *rh7*-positive neurons that express CsChrimson, and place the flies in an arena with red and blue lights on different sides, the proportion of flies on the blue side increases significantly. Thus, stimulation of *rh7* neurons causes aversion. In addition, when we exposed non-diapausing flies to pulses of blue light at night, there was a major reduction in sleep following the pulses. However, nighttime pulses of blue light did not stimulate wakefulness in the *rh7* mutant. Thus, our data support the model that flies prefer to sleep in the shade because blue light stimulation of *rh7* neurons causes both aversive behavior and arousal. Our results are consistent with a previous finding that *rh7* mutants fail to avoid blue light during the day (54). This study tracked the absolute location of the flies over the span of a day, and did not distinguish whether or not a fly was asleep. Here, we demonstrate that daytime blue-light avoidance is primarily driven by a preference for where to rest, and this behavior depends on Rh7.

The question arises as to why diapausing flies lack the Rh7-mediated preference for sleeping in the shade during the day, as occurs in non-diapausing flies. A plausible explanation for this behavior is that their increased sleep pressure overwhelms their aversion to sleeping under blue light. Flies in diapause display increased activity of R2 neurons and expression of Bruchpilot, both of which indicate an increased sleep drive. Partially alleviating this sleep drive, by blocking the activity of R2 neurons, increased the preference for shaded sleep during diapause. Conversely, pharmacologically increasing sleep drive in flies at a warm temperature decreased this preference. These data support the model that the effects of cool temperature on sleep in diapausing flies are active responses. We propose that flies in a diapausing environment are relatively indifferent to sleeping in shade owing to elevated sleep pressure, rather than an inability to sense blue light.

### Daytime sleep behavior in diapausing flies resembles narcolepsy

Our work reveals that diapausing flies do not just sleep more than non-diapausing flies. Rather, they enter a far deeper sleep state than non-diapausing flies, as their sleep is characterized by a much higher arousal threshold, which is one of the defining characteristics of a deep sleep state. Moreover, this deep sleep state occurs during both the day and night. This finding is reminiscent of hibernating animals, which enter a deeper sleep state than during standard sleep when they are not hibernating.

There are several other observations that highlight that diapausing flies enter a deep sleep state distinct from non-diapausing flies. The increase in the arousal threshold in diapausing flies is not only higher, it is profound even after very short periods of inactivity. Remarkably, after just 15–60 seconds of inactivity, diapausing flies exhibit an arousal threshold that is greater than the highest average threshold ever exhibited by non-diapausing flies, which occurs after 60–90 minutes. Moreover, the sleep latency of diapausing flies remains very low at all times during the day and night. This is very different from non-diapausing flies, in which the sleep latency is short only at night, and for a brief period during their siesta. Even during these periods, the sleep latency of non-diapausing flies is not as short as in diapausing flies. The combination of these findings demonstrate that diapausing flies enter a persistent deep sleep state that is not observed in non-diapausing flies. We suggest that diapausing flies display one of the key features of narcolepsy–excessive daytime sleep, since if they are awake for as little as 15 seconds during the day, they quickly return to a deep sleep-state in which they are hard to rouse. Moreover, as is common in narcoleptic animals, the circadian locomotor pattern of diapausing flies is altered dramatically.

### Implications for understanding diapause

Classically-studied diapause traits include slow growth and development, reproductive arrest, altered metabolism, increased stress resistance, and lifespan extension (2, 8). Less studied have been the effects of diapause-inducing conditions on behavior. At first glance, it seems surprising that temperature, rather than photoperiod, alters the circadian rhythm so profoundly in flies in diapause-inducing conditions. Yet, reproductive dormancy in *Drosophila melanogaster* is also known to depend more strongly on cool temperatures than on short day length (17). By contrast in species with a strong (12) photoperiodic influence on diapause, such as high latitude *Drosophila* strains, there is a stronger dependence on photoperiod(9, 17).

Some investigators define diapause as a photoperiod-dependent reproductive arrest and prefer to call temperature-dependent effects “dormancy.” However, the effect of photoperiod can alternatively be interpreted as a shift in the temperature at which >50% of females cease egg development (i.e., the critical temperature) (9). Temperature may be the more salient information species because temperature rather than photoperiod directly affects growth rates and reproductive success.

The influence of photoperiod seems to be selected for at high latitudes (9). Both low and high latitude strains of *Drosophila montana* arrest reproduction at low temperatures, whereas only high latitude strains show a strong dependence on photoperiod. An advantage of responding to photoperiod rather than directly to environmental temperature is that animals can anticipate and prepare in advance for harsh conditions using photoperiod, which is less susceptible to unseasonal variation. However a potential disadvantage to relying on photoperiod is that during periods of climate change, photoperiod could become a poor predictor of temperature, which is the feature with a direct impact on species survival. An advantage of responding directly to temperature might be conferred closer to the equator where variations in photoperiod are less extreme.

The effects of cool temperatures on sleep and activity that we report here are also independent of the hormone JH, which is a key regulator of diapause that promotes vitellogenesis (1, 8). We previously found that cool temperatures also arrest germline stem cell division independently of JH (13). Together with the current findings of JH-independent effects on circadian rhythm and sleep characteristics (*SI Appendix,* Fig. S2), our results suggest that cool temperatures exert multiple effects on fly physiology, behavior, and ovarian development that together represent a holistic, adaptive response.

## Materials and Methods

### Drosophila stocks and maintenance

Flies stocks and crosses used in this work were reared and maintained in temperature (25 °C) and humidity (60%) controlled incubators (Darwin Chambers) under a 12 hr light (∼750 lux):12 hr dark cycles on standard fly food (molasses, yeast, and cornmeal). Prior to the start of experiments testing behavior, flies were allowed to acclimatize to their arena for ≥16 hours. The following fly stocks were used: Canton S (BDSC#64349); *w^1118^*(BDSC#5905); *trpl^MB10553^*;*trp^MB03672^* (C. Montell lab); *per^01^*,*w^1118^* (BDSC#80917); *w^1118^,norpA^P24^* (C. Montell lab); *w\**;*clk^out^*(BDSC#56754); *UAS*-*ClkΔ* (BDSC#36319); *R30G03-Gal4* (BDSC#49646); *CaLexA* (*lexAOP-*GFP;;*UAS-LexA::VP16::NFAT*/TM6B) (BDSC#66542); *clk.856*-*Gal4* (BDSC#93198); *UAS-TNT* (BDSC#28838); *GMR-hid* (C. Montell lab); *lexAOP-CsChrimson* (BDSC#55137).

### Generation of *rh7^LexA^* flies

To generate the *rh7^LexA^* allele we used CRISPR/Cas9 to insert the *LexA* gene in frame in place of the first 179 base pairs (bp) of the second exon, following the endogenous translation start site. The construct also included the *mini white* (*w^+^*) genes. The donor plasmid was pBPLexA::p65Uw (Addgene). The two homology arms were amplified from genomic DNA prepared from *w^1118^* using the following primers: forward primer for 5′ arm: GAAAAGTGCCACCTGCTCACATGCGAAGGGGGAAT; reverse primer for 5′ arm: CATTTTGATTGCTAGGTGGTACTTGGCCAAATATTTACGAGC; forward primer for 3′ arm: GACAAGCCGAACATAGAGAATCCCACAAATGAATTTCTGCAAG, reverse primer for 3′ arm: CTAGGCGCGCCCATAGTGAATGTAGGCAAGGACCACG. The 5′ arm was inserted between the AatII and NheI sites, and the 3’ arm was inserted into the NdeI site. The 5’ arm included a modified Kozak sequence: ACCAC. The guide-RNA targeting sequence was: GATGGCCTCCATGACTGACT.

### Drosophila Activity Monitor assay

To record flies’ circadian activity profiles at various temperatures and light:dark cycles, 5–10 day-old mated female Canton S flies were recorded. Flies were individually loaded into clear glass tubes with a sucrose food source (5% sucrose + 1.5% agar) on one end and a cotton plug on the other. Flies were allowed to acclimatize for >24 hours prior to the start of a 4-day-long recording. Their activity was logged on a computer using the *Drosophila* Activity Monitor system (DAM; Trikinetics; (19)) and plotted in 30-minute bins using the Rethomics R software package (65). Moving/30 minutes represents the proportion of time in which there were beam breaks per 30 minutes. In these experiments, flies were housed under white LED lights (Minger #H6150) at ∼1700 lux. Dead or sick flies (i.e. flies with <14 beam breaks/day) were censored from the analysis.

### Behavioral arena for activity/sleep monitoring and shade preference

Given the limited spatial resolution of the DAM assay, we designed a custom behavioral arena to video track the movement of flies throughout the day, which also served as the platform used to assess flies’ daily preference for shade, preference for red vs. blue light, and startle response to vibration or light stimuli. This 102 mm x 119 mm x ∼7 mm arena individually-housed 30 flies, each in a rectangular enclosure (44 mm x 6 mm x 6 mm) with ad libitum access to a 5% sucrose in 1.5% agar. The face of the arena contained grooves to fit a clear acrylic sheet (McMaster #8560K171) that prevented the flies from escaping. Flies were anesthetized with CO_2_ in order to load them into their enclosure, and then allowed ∼24 hours to recover prior to the start of the recording. For experiments testing flies’ preference for shade, we covered one half of the arena with a 0.9 neutral density filter, which blocked visible light but was nearly transparent to our IR light source. We used a similar approach in experiments testing the preference for red vs. blue, where one half of the arena was covered with one of the light filters. In order to visualize flies at night, as well through a light filter, the arena was backlit with an 850 nm near-IR LED light (Waveform #7031) and recorded with an ELP 2MP webcam affixed with an IR-pass filter (Heliopan 850 nm). A CCS100 Spectrometer (Thorlabs) was used to measure the relative light intensity in our various experimental conditions (*SI Appendix*, Fig. S1*A* and Fig. S4*A*). The STL file for the behavioral arena can be found at the following URL: https://github.com/Craig-Montell-Lab/Meyerhof-et-al.-2024-/tree/main/SunSeekerPrintFile

To analyze movement and sleep in flies in our arena, we wrote custom Matlab scripts (https://github.com/Craig-Montell-Lab/Meyerhof-et-al.-2024-). Dead or sick flies were censored from the analysis and were defined as any fly with a bout of immobility that lasted >6 hours. Data are displayed in 30-minute bins unless otherwise indicated, and the preference index for shaded sleep was defined as:

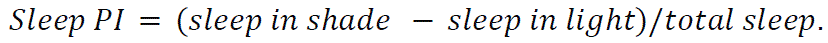

### Video tracking

To track the position of flies in the aforementioned arena we wrote custom scripts in Matlab 2021a (Mathworks), which is available at https://github.com/Craig-Montell-Lab/Meyerhof-et-al.-2024-. Flies were recorded at ∼2 frames per second in grayscale at a resolution of 1080 x 19020 from a webcam placed 45 cm away from the arena. Fly positions were recorded in real time via a movement-based tracking algorithm. To identify foreground pixels (i.e., pixels corresponding to moving flies) over multi-day experiments, we implemented a dynamic background model. The background model was created by taking the average pixel intensity at each pixel location from 30 consecutive frames. Then, a new frame was read in and the absolute difference was taken between it and the background model. Foreground pixels were defined as those greater than an empirically determined threshold (6 pixels):

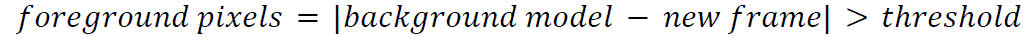

Following this step, the oldest frame in the background model was replaced by the new frame and the process repeated. To reduce the chances of misassigning background pixels as foreground, we only considered contiguous foreground pixel regions (blobs) >10 pixels^2^ and <500 pixels^2^ that occurred within a fly enclosure. An example of the tracking is given in Supplementary Video 1.

### Vibration sleep arousal threshold

To test the sleep depth of flies at 25 °C and 10 °C we housed them in our behavioral arena (under identical conditions to what has been described above) and subjected them to five gradually-increasing vibration stimuli once every two hours over the span of a day (Fig. 3*B* and *SI Appendix*, Fig. S3A). This was accomplished by applying 1–5 V, via pulse-width modulation from an Arduino Uno (Elgoo) microcontroller, to a Mosfet transistor (WeiMeet) that controlled four vibrating motors wired in parallel and placed at each corner of the rectangular arena (Fig 3*A*).

We used a triple-axis accelerometer (ADXL326; Elgoo) wired to an Arduino Uno (Elgoo) to measure the *g* force on the arena as voltage was applied to the vibrating motors (*SI Appendix*, Fig. S3 *A* and *B*).

### Blue light sleep arousal

To test the wake-promoting effects of blue light (Fig. 5 *D*–*I*), we subjected flies to five 3 second light pulses with a 10 second interpulse delay, once every two hours. We used a blue (445 nm) LED light strip (85 lux; American Bright Optoelectronics Corp.) that was controlled via an Arduino Uno (Elgoo), which received commands from a custom Matlab script (https://github.com/Craig-Montell-Lab/Meyerhof-et-al.-2024-/tree/main/SunSeeker_Ligh tPulse) that was integrated into our tracking program.

### Immunohistochemistry and confocal microscopy

All immunostaining was performed using whole mount preparations of adult brains. Whole flies were fixed in 4% paraformaldehyde (Electron Microscopy Sciences) and 0.3% Triton X-1000 in 1x PBS for 1 hour at 4 °C. Samples were washed 2 times in 0.3% PBST (0.3% Triton X-1000 in 1x PBS). Then the brains were dissected into fresh, chilled PBST, blocked in 0.3% PBST containing 5% normal goat serum (NGS) for 1 hour at room temperature, and incubated with primary antibodies in 0.3% PBST + 5% NGS for 24 hours at 4 °C. Samples were then subjected to three 30-minute washes, and incubated in secondary antibodies in 0.3% PBST + 5% NGS overnight (∼16 hours) at 4°C. After three additional 30 minute washes, the brains were mounted on glass slides with VECTASHIELD anti-fade mounting media (Vector Labs, catalog. H-1200). Primary antibodies: anti-GFP (chicken, Invitrogen, A10262, 1:1000 for CaLexA expression comparison), anti-nc82 (mouse, Developmental Studies Hybridoma bank, nc82 concentrate, 1:50 for BRP expression comparison, else 1:250 as structural stain) Secondary antibodies: Alexa Fluor 488 conjugated goat anti-chicken (1:1000, Invitrogen, A11039), Alexa Fluor 633 conjugated goat anti-mouse (1:1000, Invitrogen, A21050).

### Acquisition and quantification of images

All images were acquired using a Zeiss LSM 900 confocal microscope at 20x magnification (PApo 20x/0.8 objective) at a resolution of 3064 x 3064 pixels. Within each experiment, images were acquired using identical laser power, gain, Z-stack depth (1µm), and scan speed using the CO-2Y setting (Zen Blue). Unprocessed images were exported from Zen Blue as.czi files and imported to FIJI (ImageJ). To measure signal intensity, the background signal of each optical slice in a stack was subtracted (except in the case of BRP experiments), a maximum intensity projection was generated, and the mean pixel intensity of each outlined ROI (whole brain or EB) was calculated.

### Optogenetics

We used the channelrhodopsin csChrimson to activate *rh7* neurons in order to test their effect on color/sleep preference. *rh7^LexA^* virgin females were crossed to *lexAOP-*csChrimson males on standard fly food with or without the addition of 1 mM all-*trans* retinal (Sigma;R2500). After four days, the parents were removed from the fly vial and the progeny were shifted to constant darkness to complete development. After eclosion and mating, female *rh7>CsChrimson* flies were transferred to our behavioral arena which contained a red– and blue-light filter (Neewer.com).

### Testing effect of gaboxadol on shaded sleep preference

To test the effect of sleep pressure on shaded sleep preference, we loaded flies into our behavioral arena with a sucrose food source (5% sucrose + 1.5% agar) that contained either 0.1 or 1 mg/mL of gaboxadol (Cayman; 16355). Mated female Canton S flies were aspirated via mouth pipette into the arena at ZT 8 the day prior to the start of the recording, where they had ad libitum access to gaboxadol-laced food throughout the experiment.

### Quantification and Statistical Analysis

The sample sizes for each test are provided in the figure legends. “n” denotes the number of flies examined. All statistical tests were performed in R (version 4.1.0). Based on our experience and common practices in this field, we used a sample size of n ≥ 20 flies for experiments monitoring circadian locomotor and sleep patterns, with the exact sample size displayed in the figure legend. Non-parametric Aligned-Rank ANOVA was performed using the “ARTool” library. Parametric ANOVA was performed using the “stats” R package. Plotting was performed using either the “ggplot2” R library or Matlab 2021a. Error bars display SEMs unless otherwise indicated in the figure legend. We set the significance level, α = 0.05. Asterisks indicate statistical significance: *P < 0.05, **P < 0.01, and ***P < 0.001.

## Data Availability

All study data are included in the article, and in the SI Appendix.

## Author Contributions

G.T.M., A.E.B., S.E., D.J.M, and C.M., designed experiments. G.T.M., A.E.B., and S.E. performed experiments. G.T.M., A.E.B., and S.E. analyzed data. G.T.M. designed and edited figures. G.T.M, S.E., D.J.M., and C.M. wrote the first draft of the manuscript. G.T.M., S.E., A.E.B., D.J.M, and C.M. provided edits and critical feedback related to this work.

## Classification

Biological sciences and neuroscience

## Acknowledgments

We thank Nicole Y. Leung for generating the *rh7^LexA^* line, and Nicolas A. Debeaubien for his assistance in designing the shade-preference behavioral arena, as well as creating the images shown in Fig. 2*A* and 3*A*. This work was supported by a grant to DJM from the National Institute on Aging (R01AG36907) and by grants to CM from the National Institute of Allergy and Infectious Disease (R01AI169386) and the National Institute on Deafness and Other Communication Disorders (R01DC016278). GM was supported in part by a predoctoral fellowship from the National Eye Institute (F31EY033179), and AEB was supported in part by a predoctoral fellowship from the National Institute on Deafness and Other Communication Disorders (F31DC021112).

## Figure legends

**Fig. S1.**
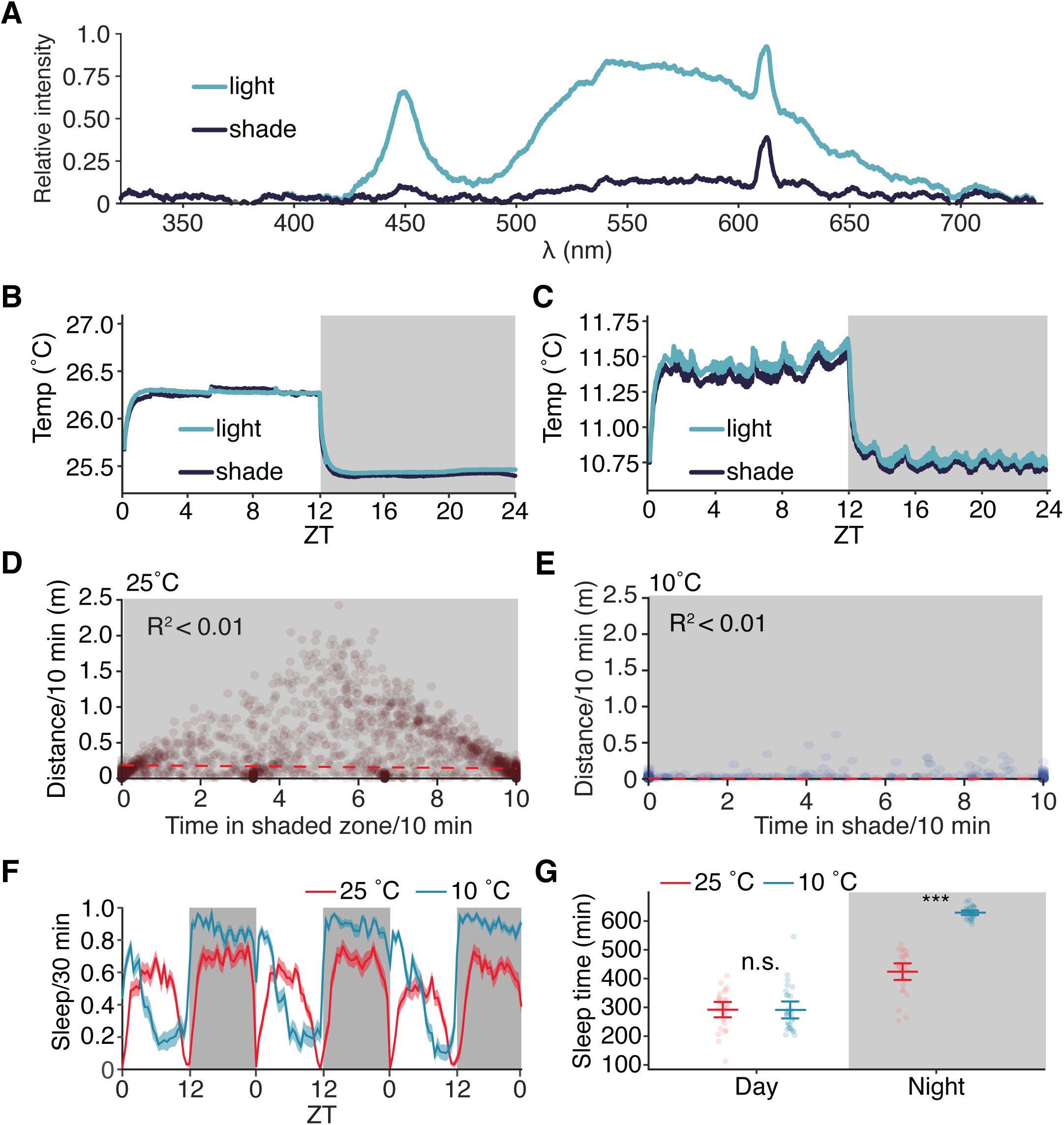
Light intensity, distance vs. shade correlation, and sleep of flies in diapause. (*A*) Relative light intensity of shaded and unshaded zone in arena. (*B* and *C*) Temperature recording from behavioral arena (Fig. 2*A*) from shaded and unshaded zones over the span of a day when the arena was housed in an incubator set to 25°C (A) or 10 °C (B). (*D*) Correlation between distance/10 minutes and time spent in the shade/10 minutes from flies at 25 °C. Each dot represents a 10 minute bin from the night time in a three-day long recording. Red dashed line represents the linear best fit line. (*E*) Correlation between distance/10 minutes and time spent in the shade/10 minutes from flies at 10 °C. Each dot represents a 10 minute bin from the night time in a three-day long recording. Red dashed line represents the linear best fit line. (*F*) Three-day sleep profile of flies housed at 25 °C (red) or 10 °C (blue). Shading represents SEM. (*G*) Quantification of average sleep time during three-day long recording, separated by day and night. Error bars indicate the 95% confidence interval around the mean. Data were analyzed first by Aligned Rank Transform Two-Way ANOVA, examining factors of time, temperature, and time:temperature interaction. Individual differences between groups were analyzed by Aligned Rank Transform Contrast, with Bonferroni multiple testing correction. *D*–*G*: n = 27–29 flies/temperature. \*\*\**P*<0.001.

**Fig. S2.**
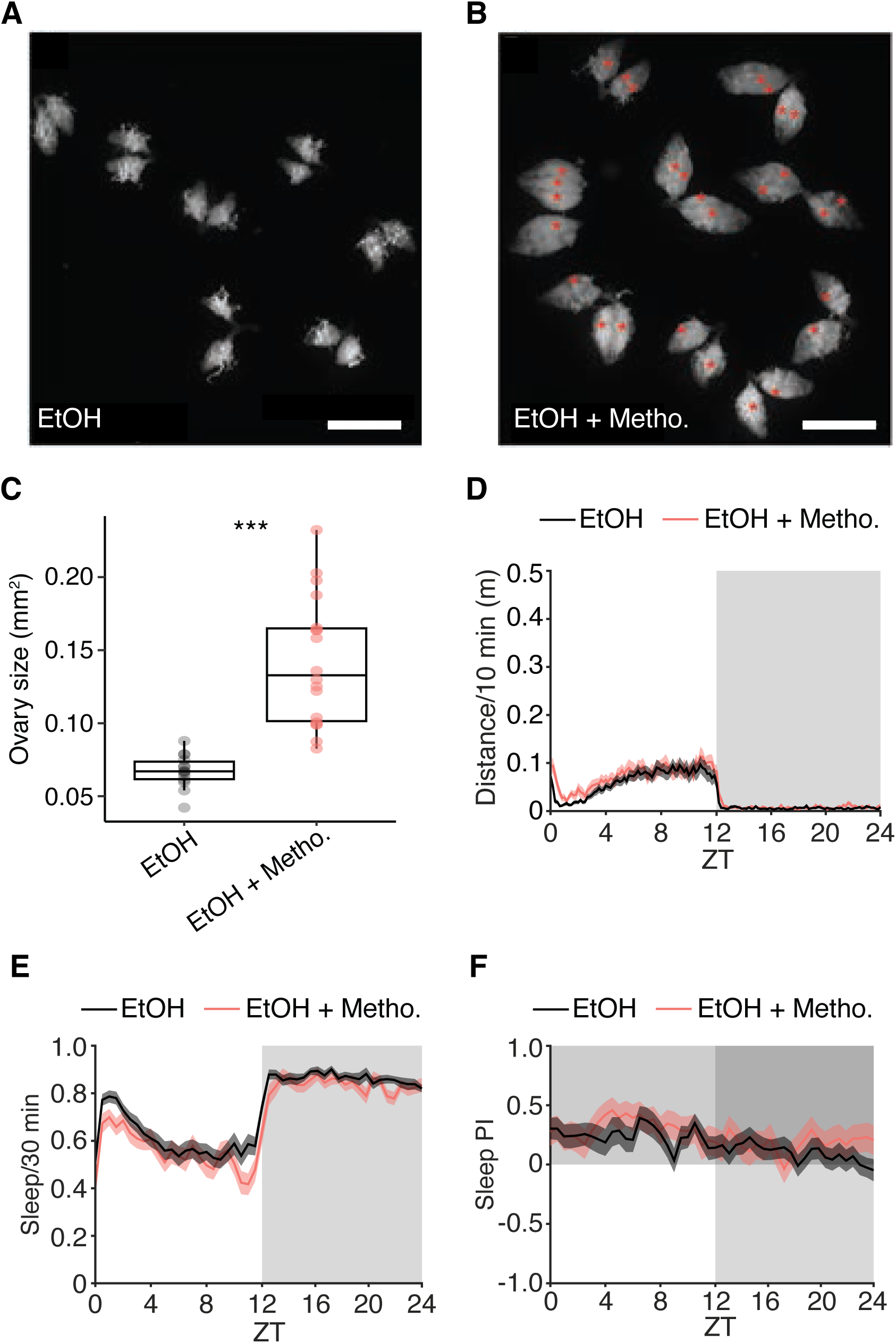
Methoprene treatment reverses ovarian arrest in diapause but fails to modify circadian behaviors. (*A*) Darkfield micrograph of the whole ovary from flies treated with ethanol (drug vehicle control). Scale bar is 1 mm. (*B*) Darkfield micrograph of whole ovaries from flies treated with ethanol and 100 µg of methoprene. Scale bar is 1 mm. Asterisks label egg chambers ≥ stage 12. (*C*) Quantification of ovary size as area (mm^2^), performed using Fiji software. Box plot represents median, 25th percentile, and 75th percentile, with whiskers extending to either to the min/max value or 1.5 x the interquartile range. n=14–18 ovaries per condition. Groups compared via Wilcoxon rank sum test. \*\*\**P<0.001.* (*D*) Circadian activity profile of vehicle-treated (black) or methoprene-treated (red) flies at 10 °C. (*E*) Sleep profile of vehicle-treated (black) or methoprene-treated (red) flies at 10 °C. (*F*) Shaded sleep PI profile of vehicle-treated (black) or methoprene-treated (red) flies at 10 °C

**Fig. S3.**
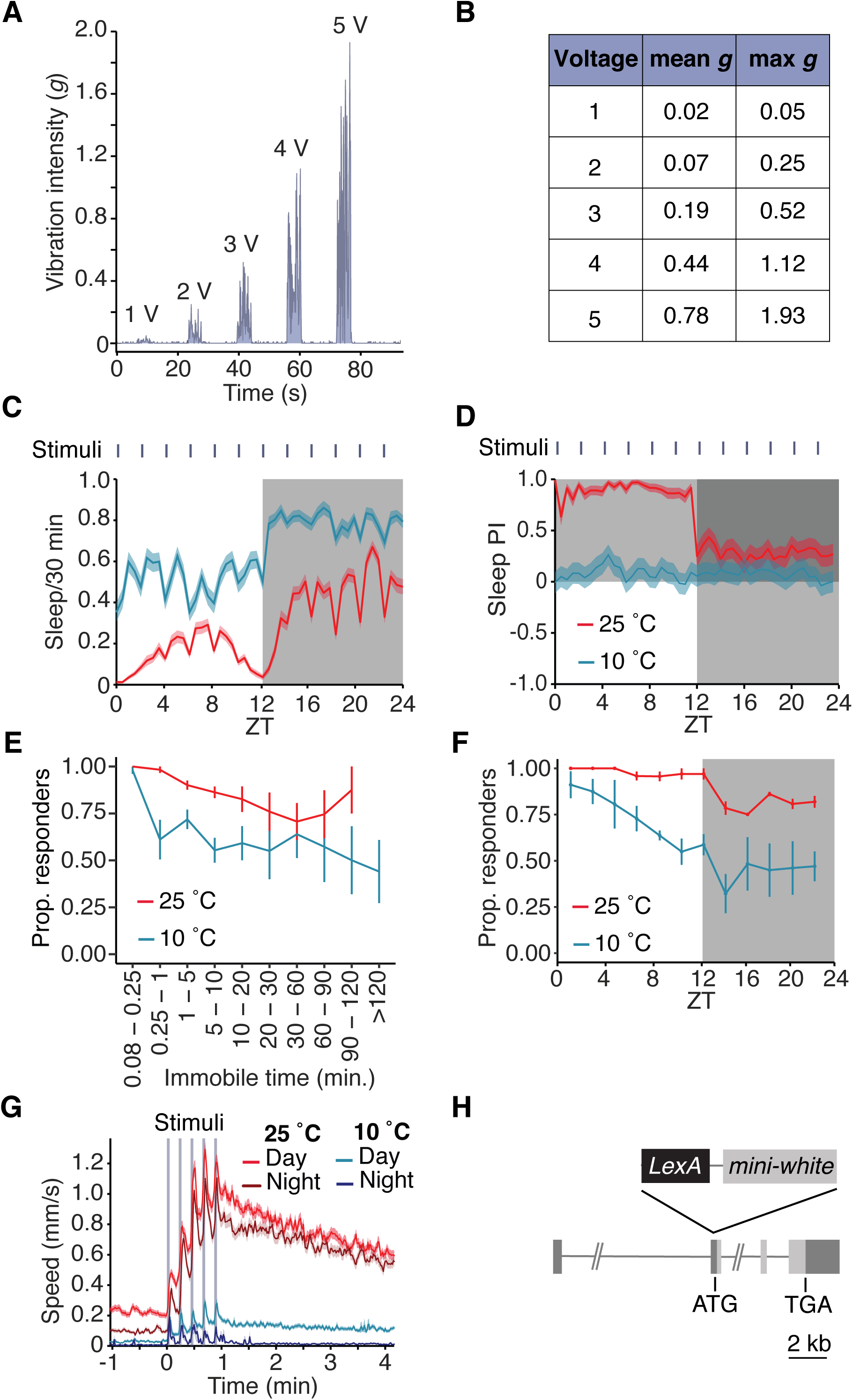
Vibration intensity and arousal threshold of flies in diapause. (*A*) Vibration intensity (*g* force) of the behavioral arena (Fig. 3*A*) when 1–5V are delivered to the vibrating motors via pulse-width modulation from an Arduino microcontroller. (*B*) Table summarizing the vibration intensity on the arena when 1–5 volts is delivered to the four vibrating motors (Fig. 3*A*) flanking the behavioral arena. (*C*) Average sleep profile of flies at 25°C (red) or 10°C (blue) during arousal threshold experiments. “Stimuli” denotes the start of the stimulus train, which was delivered to flies once every two hours. Shading represents SEM. (*D*) Average sleep preference index of flies at 25°C (red) or 10°C (blue) during arousal threshold experiments. “Stimuli” denotes the start of the stimulus train, which was delivered to flies once every two hours. Shading represents SEM. (*E*) The proportion of flies that responded to any vibration stimulus vs. their length of immobility when maintained at 25°C or 10°C. Line denotes the group mean and error bars denote SEM. (*F*) The proportion of flies that responded to any vibration stimulus vs. the time of the day when maintained at 25°C or 10°C. We did not observe at 25 °C flies that were immoble for >120 min. Means ±SEMs. (*G*) Truncated version of plot shown in Fig. 3*H* to aid visualizing differences in walking speed between groups and in response to vibration stimuli. Plot displays average walking speed of flies in response to vibration stimuli at 25 ° or 10 °C during the day and night. Time 0 denotes the start of the stimulus train, with the vertical gray shading indicating each vibration stimulus. C–G: data recorded from 149 *w^1118^* flies over three independent experiments. (*H*) Cartoon depicting *rh7^LexA^* allele. To generate *rh7^LexA^* 179 bp were deleted after the translation start site in the second exon, and in its place a *LexA* and *mini-white* were inserted in frame.

**Fig. S4.**
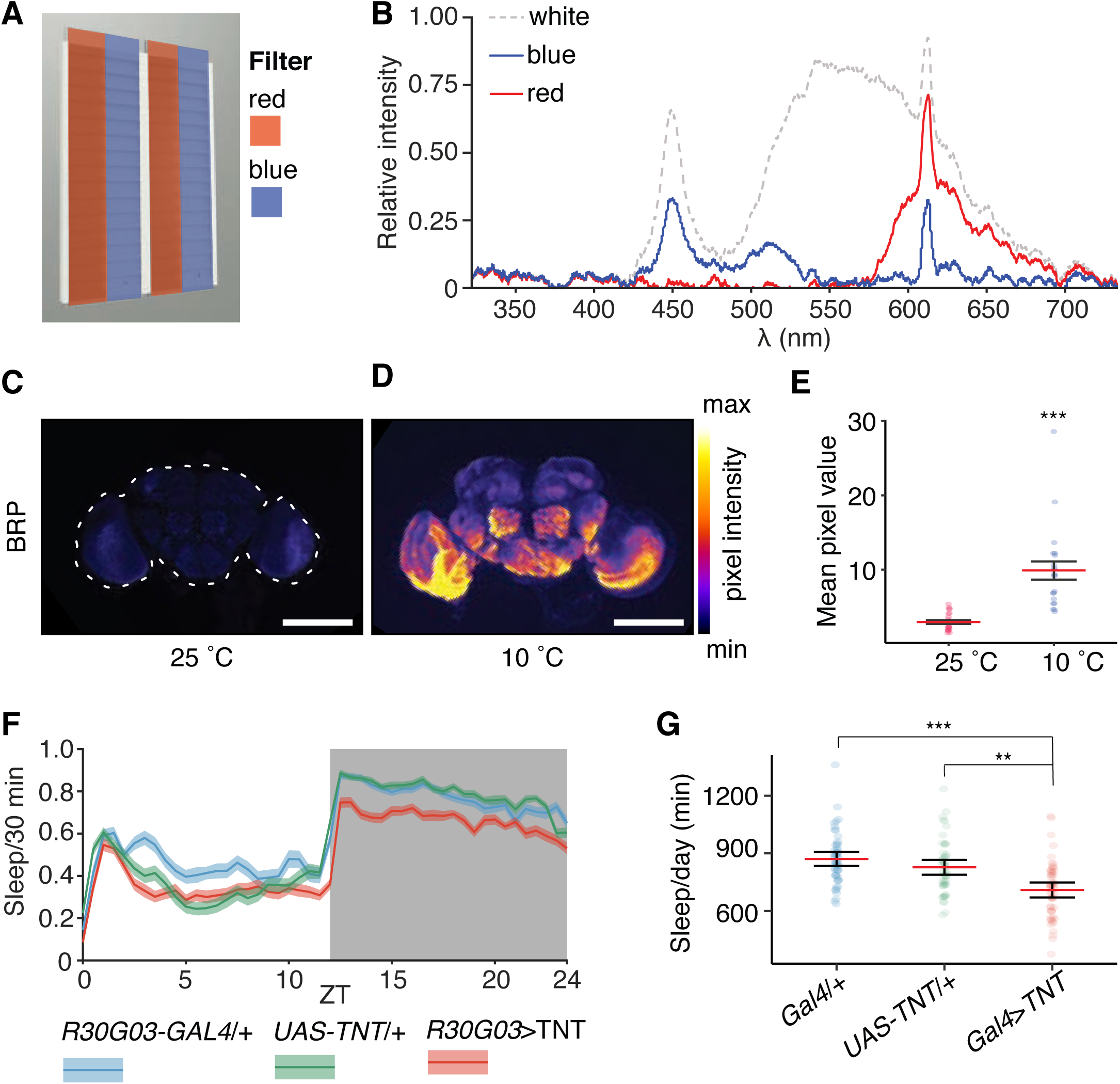
Red versus blue arena, *rh7* BRP staining, and *R30G03*>TNT sleep. (*A*) Diagram of behavioral arena testing red vs. blue preference. (*B*) Relative light intensity in behavioral arena with red or blue filter. (*C*) Representative images showing BRP staining in *rh7^LexA^* flies maintained at 25 °C Scale bar indicates 100 µm. (*D*) Representative images showing BRP staining in *rh7^LexA^* flies maintained at 10°C. Outline (white dashed line) circumscribes the brain. Scale bar indicates 100 µm. (*E*) Quantification of mean pixel intensity of brains shown in B. Groups compared via Wilcoxon rank sum test. \*\*\**P*<0.001. n=21–22 brains/temperature. (*F*) Sleep profile of *R30G03*>TNT flies at 10°C. (*G*) Quantification of total sleep per day of *R30G03*>TNT shown in D. Aligned-Rank Transform ANOVA with group differences compared via Aligned Rank Transform Contrasts with Bonferroni multiple testing correction. \*\*\**P<0.001*.

